# IMiDs induce FAM83F degradation via an interaction with CK1α to attenuate Wnt signalling

**DOI:** 10.1101/2020.05.25.114660

**Authors:** Karen Dunbar, Thomas J. Macartney, Gopal P. Sapkota

## Abstract

Immunomodulatory imide drugs (IMiDs) bind CRBN, a substrate receptor of the Cul4A E3 ligase complex, enabling neo-substrate recruitment and degradation via the ubiquitin-proteasome system. Here, we report FAM83F as such a neo-substrate. We recently showed that the eight FAM83 proteins (A-H) interact with members of the serine/threonine protein kinase CK1 family, to regulate their subcellular distribution and distinct biological roles. CK1α is a well-established IMiD neo-substrate and we demonstrate here that IMiD-induced FAM83F degradation requires its association with CK1α. Despite all FAM83 proteins interacting with CK1α, no other FAM83 protein is degraded by IMiDs. FAM83F is localised to the plasma membrane, and consistent with this, IMiD treatment results in depletion of both FAM83F and CK1α levels from the plasma membrane. We have recently identified FAM83F as a mediator of the canonical Wnt signalling pathway. The IMiD-induced degradation of FAM83F attenuated Wnt signalling in colorectal cancer cells and removed CK1α from the plasma membrane, mirroring the phenotypes observed with genetic ablation of FAM83F. Intriguingly, in many cancer cell lines, IMiD-induced degradation of CK1α is only modest and incomplete. In line with this observation, the expression of FAM83G, which also binds to CK1α, appears to attenuate the IMiD-induced degradation of CK1α, suggesting a protective role for FAM83G on CK1α. Our findings reveal that the efficiency of target protein degradation by IMiDs, and perhaps other degraders such as PROTACs, relies on the nature of the inherent multiprotein complex in which the target protein exists. Our findings unearth opportunities for developing degraders to target specific protein complexes.

## INTRODUCTION

Thalidomide, the first immunomodulatory imide drug (IMiD), initially came to prominence as a treatment for morning sickness in the 1950s but was quickly abandoned after it became apparent that consumption of thalidomide in the first trimester of pregnancy caused foetal abnormalities, predominately manifesting as limb deformities (1). Despite these severe teratogenic effects, the mechanism of action remained elusive for several decades until it was found that IMiDs hijack the ubiquitin-proteasomal system (UPS) to facilitate protein degradation of non-native substrates, which have been termed “neo-substrates” (2). IMiDs act as a molecular glue by binding to both neo-substrates and a hydrophobic binding pocket of cereblon (CRBN), which is a substrate receptor of the Cul4A-E3 ligase complex. This brings the neo-substrates into close proximity to the Cul4A-ROC1-DDB1-CRBN E3 ligase complex (known as Cul4A^CRBN^), thereby facilitating their ubiquitylation and subsequent proteasomal degradation (2). Recently, IMiDs have been repurposed for the effective treatment of multiple myeloma with lenalidomide (3) and pomalidomide (4), two distinct derivative analogues of thalidomide, routinely used in the treatment of multiple myeloma. Their efficacy has been attributed to the induced degradation of the zinc-finger transcription factors IKZF1 and IKZF3 which have key roles in B and T cell biology (2).

Whilst the majority of identified IMiD neo-substrates appear to be zinc-finger transcription factors (2, 5, 6), lenalidomide has also been shown to induce the degradation of the serine/threonine kinase CK1α (7). Casein kinase 1 isoforms (α, α-like, δ, ε, γ1, γ2 and γ3) are a family of serine/threonine protein kinases which control many cellular processes, including Wnt signalling, circadian rhythms, calcium signalling, cell division and responses to DNA damage (8–11). Lenalidomide binds to a ß-hairpin loop in the kinase N-lobe of CK1α, bringing it into proximity of the Cul4A^CRBN^ complex to facilitate its ubiquitylation and subsequent proteasomal degradation (12). The degradation of CK1α is thought to cause the efficacy of lenalidomide in the treatment of myelodysplastic syndromes (MDS) (13). MDS are a group of blood cancers, of which a subtype are caused by deletion of chromosome 5q (del(5q)) (13). In such cancers, deletion of a region of chromosome 5q results in CK1α haploinsufficiency through loss of the *CSNK1A1* gene (7), thereby sensitizing cells against further degradation of CK1α by lenalidomide.

Historically, CK1 isoforms were thought to be monomeric, unregulated and constitutively active but there is now accumulating evidence that a family of previously uncharacterised proteins, the FAM83 proteins, act as anchors for several of the CK1 isoforms (α, α-like, δ, and ε) and can alter their subcellular localisation in response to specific stimuli (14, 15). The FAM83 family is composed of 8 members, termed FAM83A-H, which share a conserved N-terminal domain of unknown function 1669 (DUF1669), which mediates the interaction with different CK1 isoforms (14). Each member binds to different CK1 isoforms with varying specificity and affinity (14). All FAM83 proteins interact with CK1α, while FAM83A, B, E and H also interact with CK1δ and ε (14). Whilst the FAM83 family remains largely uncharacterised, roles for specific FAM83-CK1α complexes have been established in mitosis (16, 17) and canonical Wnt signalling (18–20). Given the reports of IMiD-induced degradation of CK1α, we sought to establish the effect of IMiDs on the stability of FAM83 proteins and different FAM83-CK1α complexes.

## RESULTS

### IMiDs selectively degrade FAM83F protein

Lenalidomide, which is used as a therapeutic agent in patients with del(5q) MDS, causes CK1α degradation (13). In MV4.11 cells, which are derived from B-myelomonocytic leukaemia, the thalidomide derivatives lenalidomide, pomalidomide and iberdomide led to robust degradation of IKZF1, but only lenalidomide, and to a lesser extent pomalidomide, led to partial degradation of CK1α (Fig. 1A) The efficacy of lenalidomide-induced CK1α degradation in other cell lines, including THP-1 monocytes, HCT116 colorectal cancer, A549 lung adenocarcinoma, DLD-1 colorectal cancer, PC-3 prostate cancer and HaCaT keratinocyte cell lines were more variable with the most substantial CK1α degradation observed in HCT116, DLD-1 and HaCaT cells (Fig. 1B). As the FAM83 proteins exist in complexes with CK1α (14), we sought to test the effect of the IMiDs thalidomide, lenalidomide, and pomalidomide on FAM83 protein levels in THP-1, HCT116, A549, DLD-1, PC-3 and HaCaT cell lines. Both lenalidomide and pomalidomide but not thalidomide induced a robust reduction in FAM83F protein abundance in HCT116, DLD-1 and HaCaT cells, whereas in the other tested cell lines FAM83F protein was not detectable (Fig. 1C&D). None of the IMiDs led to any detectable reduction in FAM83B, FAM83D, FAM83G or FAM83H protein abundance (Fig. 1C). Currently, no reliable antibodies exist for the detection of endogenous FAM83A, FAM83C and FAM83E proteins, limiting their assessment in this assay. A modest degradation of CK1α was observed with lenalidomide and pomalidomide treatment in HCT116, A549 and DLD-1 cells (Fig. 1C&D) whereas no consistent change in either CK1δ or CK1ε protein levels were detected following IMiD treatment (Fig. 1C). As expected, lenalidomide and pomalidomide caused robust degradation of IKZF1 in THP-1 cells, while no IKZF1 was detected in non-hematopoietic cells (Fig. 1C). ZFP91, a pomalidomide specific neo-substrate (5), was degraded upon pomalidomide treatment in all cell lines while its expression was undetectable in THP-1 cells (Fig. 1C).

**Figure 1:**
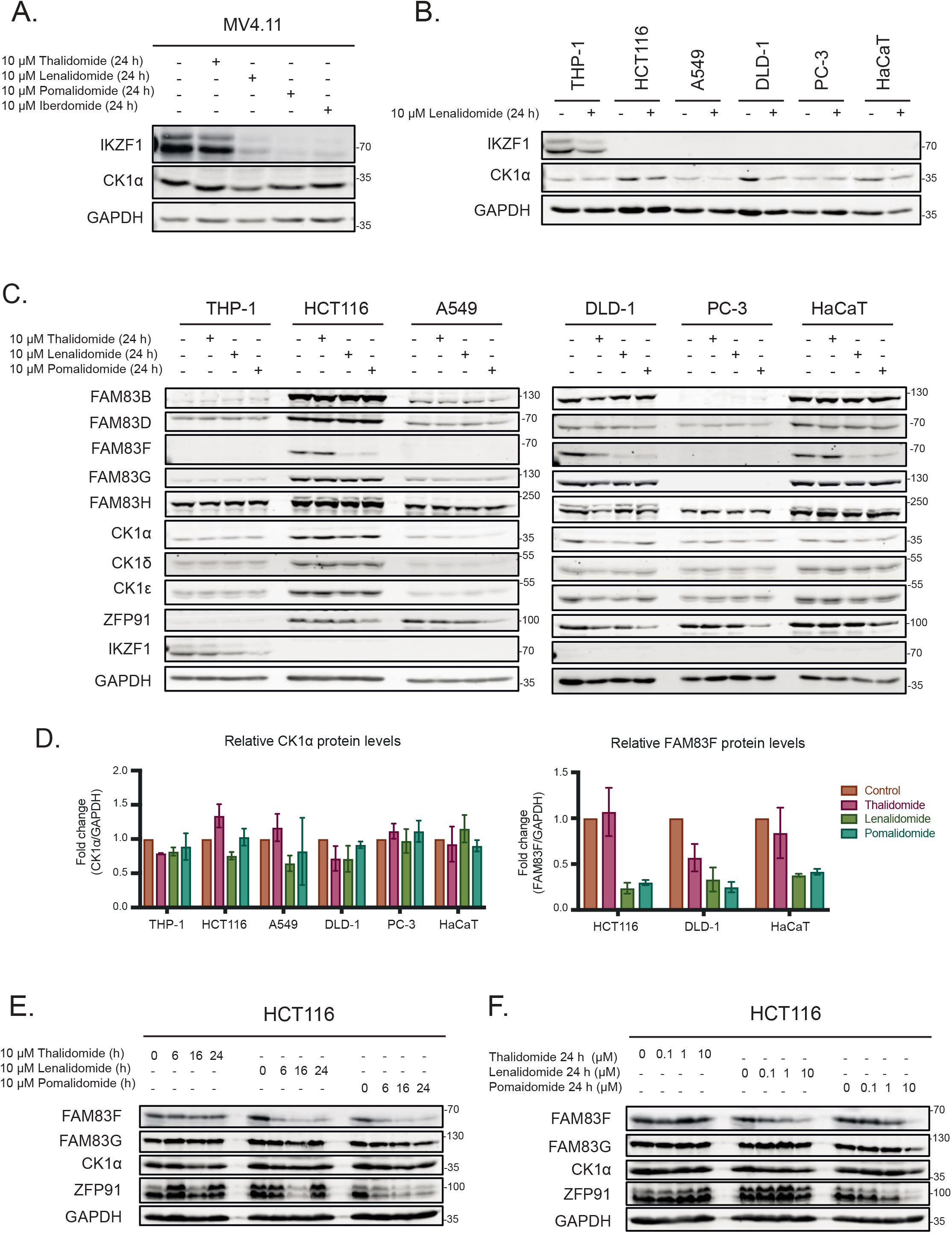
IMiDs degrade FAM83F but no other FAM83 proteins. **(A)** MV4.11 cell extracts, treated with or without the IMiD compounds (10 μM 24 h), were resolved by SDS-PAGE and subjected to Western blotting with the indicated antibodies. **(B)** THP-1, HCT116, A549, DLD-1, PC-3 and HaCaT cell extracts treated with or without 10 μM lenalidomide for 24 h were resolved by SDS-PAGE and subjected to Western blotting with the indicated antibodies. **(C)** THP-1, HCT116, A549, DLD-1, PC-3 and HaCaT cell extracts treated with or without IMiD compounds (10 μM 24 h) were resolved by SDS-PAGE and subjected to Western blotting with the indicated antibodies. **(D)** Densitometry of FAM83F and CK1α protein abundance upon treatment with the indicated IMiD compounds (10 μM 24 h). FAM83F and CK1α protein abundances were normalised to GAPDH protein abundance and represented as fold change compared to untreated cells. Data representative of two biological repeats with bar graph representing mean ± standard error. **(E)** HCT116 cell extracts, treated with 10 μM IMiD compounds for various timepoints (6, 16 and 24 h), were resolved by SDS-PAGE and subjected to Western blotting with the indicated antibodies. **(F)** HCT116 cell extracts, treated with various concentrations (0.1 μM, 1 μM and 10 μM) of IMiD compounds for 24 h, were resolved by SDS-PAGE and subjected to Western blotting with the indicated antibodies.

Further characterisation of FAM83F degradation in HCT116 and DLD-1 cells revealed a time and dose dependence of lenalidomide and pomalidomide, with optimal degradation occurring after 24 h treatments with 10 μM IMiD (Fig. 1E&F and Sup. Fig. 1A&B). Novel heterobifunctional compounds that recruit target proteins to CRBN through an IMiD moiety, such as dTAG-13, lead to target protein degradation through Cul4A^CRBN^ (21). We found that dTAG-13 was incapable of degrading CK1α, FAM83F and ZFP91 in HCT116 and DLD-1 cells, confirming the utility of certain IMiD moieties for development of bivalent protein degraders (Sup. Fig. 1C) (21).

### FAM83F and CK1α protein abundance is reduced at the plasma membrane upon IMiD treatment

Endogenous FAM83F is predominately located at the plasma membrane as observed by immunofluorescence of HCT116 cells in which a GFP tag was knocked in homozygously at the N-terminus of the *FAM83F* gene (^GFP/GFP^FAM83F cells) (Fig. 2A and Sup. Fig. 2A&3A) (20). Upon treatment of HCT116 ^GFP/GFP^FAM83F cells with pomalidomide, the GFP signal was lost from the plasma membrane (Fig. 2A). Under basal conditions, CK1α, which can interact with all eight FAM83 proteins, is distributed throughout the cell, and so we were unable to determine the effect of pomalidomide on membranous CK1α by immunofluorescence (Fig. 2A). However, when we analysed subcellular fractions of DLD-1 wild-type cells, we observed that FAM83F was predominately present in the membrane fraction, with a small proportion also observed in the nuclear fraction, whilst CK1α is present in cytoplasmic, nuclear and membrane fractions (Fig. 2B). When DLD-1 wild-type cells were treated with pomalidomide, a reduction in the levels of both FAM83F and CK1α protein was observed in the membrane fraction, while CK1α protein levels in the cytoplasmic and nuclear fractions did not change relative to untreated DLD-1 wild-type cells (Fig. 2B). FAM83F interacts with CK1α and is responsible for delivering it to the plasma membrane (20). Interestingly, the pomalidomide induced reduction of CK1α protein from the plasma membrane in DLD-1 wild-type cells was comparable to the stable reduction of membranous CK1α observed in FAM83F-knockout (FAM83F^-/-^) DLD-1 cells, generated using CRISPR/Cas9 genome editing, in the absence of pomalidomide treatment (Fig. 2B and Sup. Fig. 2A&3B). In the absence of FAM83F, pomalidomide treatment did not induce any detectable degradation of CK1α from the membrane fraction in DLD-1 FAM83F^-/-^ cells, confirming that the IMiD-induced loss of CK1α protein from the plasma membrane is mediated by FAM83F.

**Figure 2:**
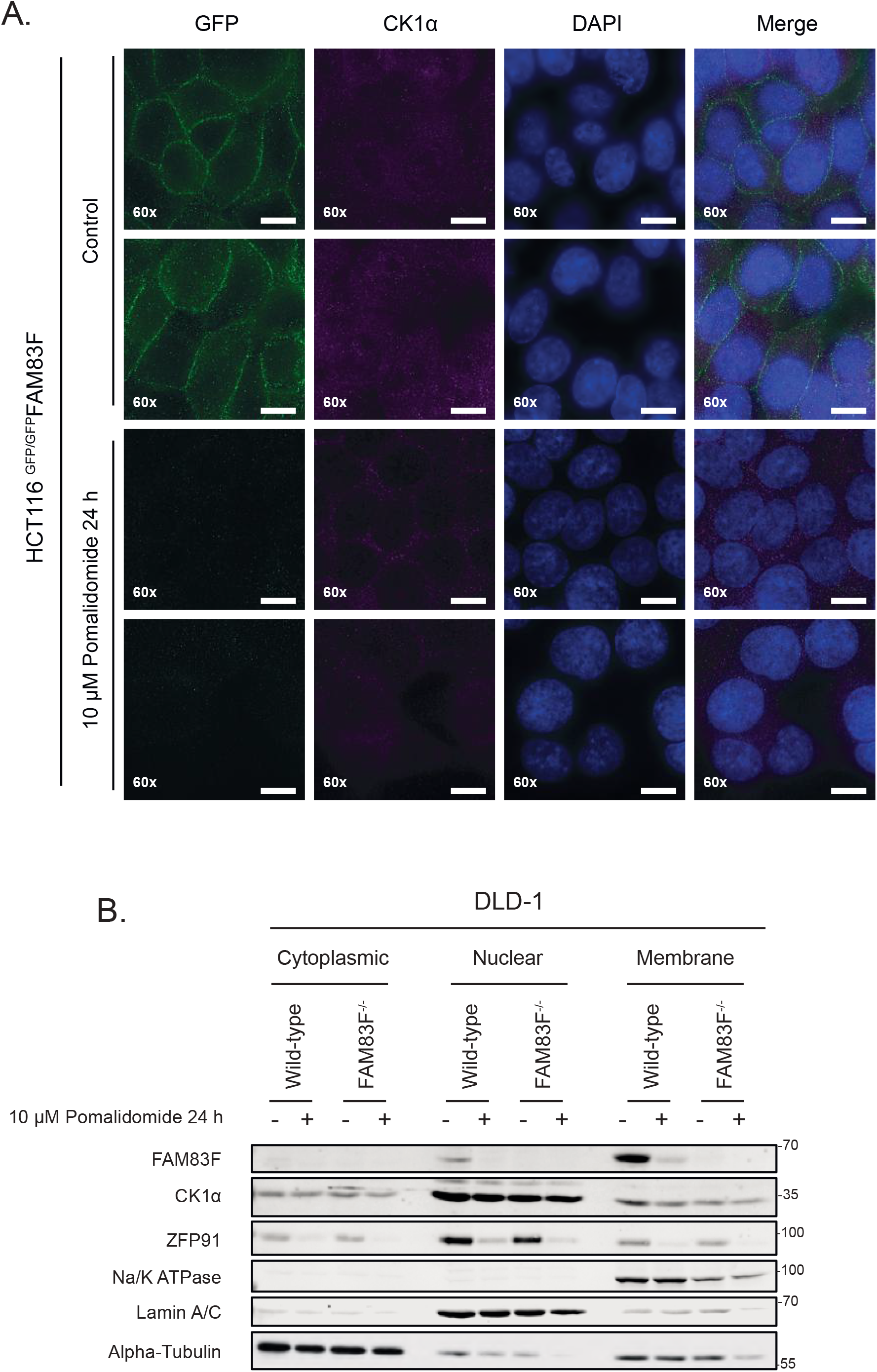
FAM83F and CK1α protein abundance is reduced at the plasma membrane upon IMiD treatment. **(A)** Widefield immunofluorescence microscopy of HCT116 ^GFP/GFP^FAM83F knock-in cells, treated with or without 10 μM pomalidomide for 24 h, stained with antibodies recognising GFP (far left panels, green), and CK1α (second row of panels from left, magenta) as well as DAPI (third row of panels from left, blue). Overlay of GFP, CK1α and DAPI images as a merged image is shown on the right. Immunofluorescence images were captured with a 60x objective. Scale bar represents 10 μm. Two representative images for each staining are shown. **(B)** Specific subcellular fractions from cytoplasmic, nuclear and membrane compartments from DLD-1 wild-type and FAM83F^-/-^ cells treated with or without 10 μM pomalidomide for 24 h, were resolved by SDS-PAGE and subjected to Western blotting with the indicated antibodies. Specificity of cytoplasmic, nuclear and membrane fractions were determined with Western blotting with compartment specific antibodies: alpha-tubulin (cytoplasmic), Lamin A/C (nuclear) and Na/K ATPase (membrane).

### IMiD-induced degradation of FAM83F requires interaction with CK1α

The robust degradation of FAM83F by lenalidomide and pomalidomide in several cancer cell lines prompted us to explore whether this degradation was mediated through the association of FAM83F with CK1α. Several conserved residues within the DUF1669 domains of FAM83 proteins have been identified as critical mediators of the FAM83-CK1 interaction (14). For FAM83F, mutation of two phenylalanine residues at positions 284 and 288 to alanine would be predicted to abolish association with CK1α (20). As expected, FAM83F co-precipitated with CK1α in CK1α immunoprecipitates (IPs) from HCT116 wild-type cells but not from FAM83F-knockout (FAM83F^-/-^) cells generated using CRISPR/Cas9 (Fig. 3A and Sup. Fig. 2A&3B). FAM83F also co-precipitated with CK1α in CK1α IPs from HCT116 FAM83F^-/-^ cells that were stably rescued with FAM83F^WT^ but not from HCT116 FAM83F^-/-^ cells rescued with FAM83F^F284/288A^ mutant (Fig. 3A). The lenalidomide and pomalidomide-induced degradation of FAM83F was only evident in HCT116 wild-type and HCT116 FAM83F^-/-^ cells rescued with FAM83F^WT^, but not in HCT116 FAM83F^-/-^ cells or in HCT116 FAM83F^-/-^ cells rescued with FAM83F^F284/288A^ mutant (Fig. 3B&C). Overall, these observations suggest that, the interaction of FAM83F with CK1α is required for the IMiD-induced degradation of FAM83F (Fig. 3B&C).

**Figure 3:**
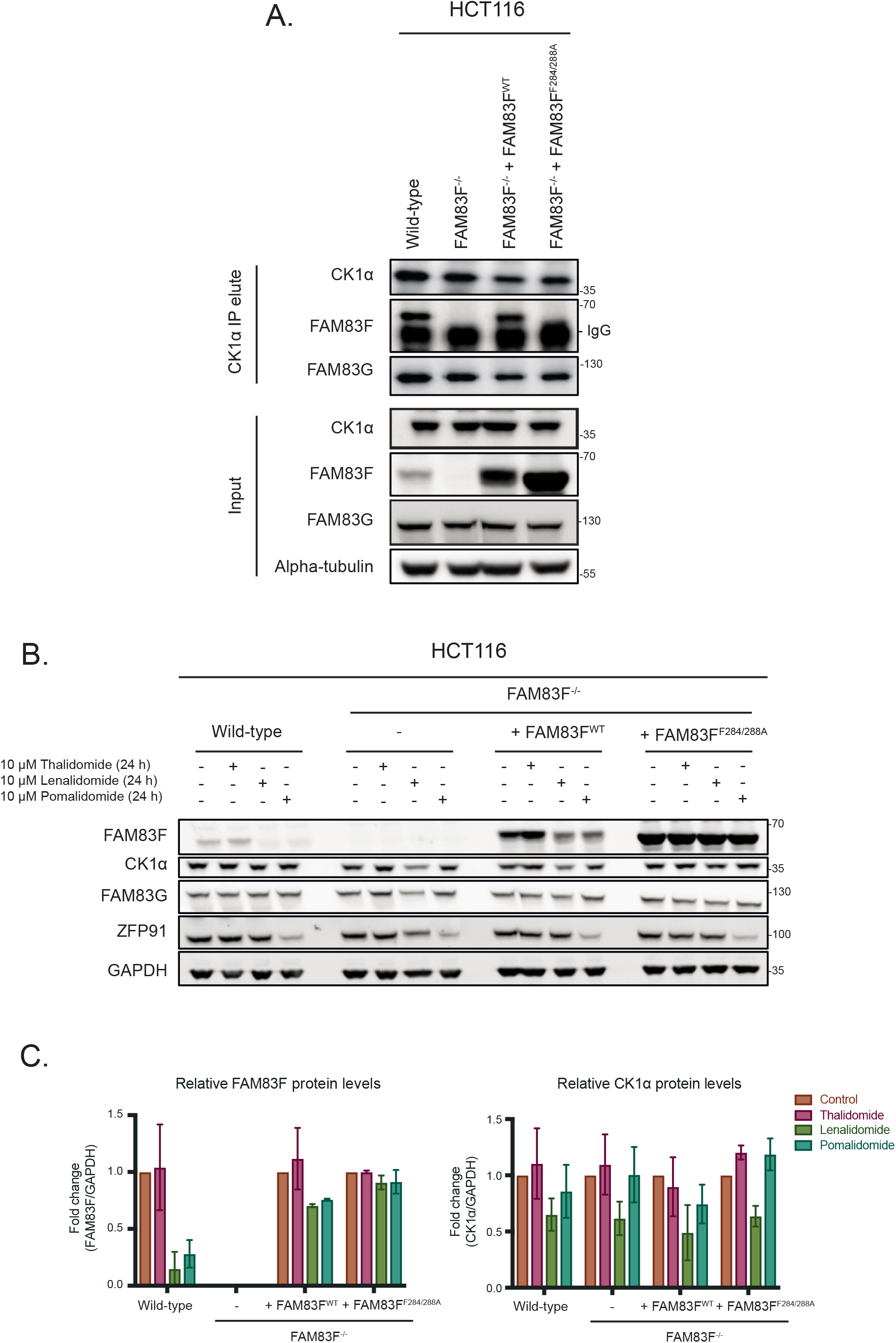
IMiD-induced degradation of FAM83F requires interaction with CK1α. **(A)** HCT116 wild-type, HCT116 FAM83F^-/-^, HCT116 FAM83F^-/-^ rescued with FAM83F^WT^ and HCT116 FAM83F^-/-^ rescued with FAM83F^F284288A^ cell extracts were subjected to immunoprecipitation (IP) with anti-CK1α antibody. Input extracts and CK1α IP elutes were resolved by SDS-PAGE and subjected to Western blotting with the indicated antibodies. **(B)** As in (A), except the cells were treated with IMiD compounds (10 μM 24 h) as indicated and then were resolved by SDS-PAGE and subjected to Western blotting with the indicated antibodies. **(C)** Densitometry of FAM83F and CK1α protein abundance upon treatment with IMiD compounds (10 μM 24 h) from (B). FAM83F and CK1α protein abundance was normalised to GAPDH protein abundance and represented as fold change compared to untreated cells. Data are representative of two biological replicates with bar graph representing mean ± standard error.

### IMiD-induced FAM83F degradation is mediated via the proteasome and is dependent on cereblon

IMiD-induced degradation requires the Cul4A^CRBN^ E3 ligase complex for the ubiquitylation of neo-substrates and subsequent proteasomal degradation. Activation of Cul E3 ligases requires NEDDylation of Cullin subunits. Thus, Cul E3 ligase activity can be blocked by inhibiting the catalytic activity of the NEDD8-activating enzyme with the small molecule inhibitor MLN4924 (22). Treatment of cells with MLN4924 prevented lenalidomide and pomalidomide-induced degradation of FAM83F in both DLD-1 and HCT116 cells, indicating the requirement of a Cul E3 ligase for degradation (Fig. 4A). Inhibition of the proteasome with bortezomib, which leads to the accumulation of mono- and poly-ubiquitylated proteins in cell extracts, also prevented lenalidomide and pomalidomide-induced FAM83F degradation, indicating that the reduction in FAM83F protein is mediated by the proteasome (Fig. 4A). To ascertain whether the IMiD-induced degradation of FAM83F was dependent on CRBN, we knocked out CRBN from DLD-1 cells using CRISPR/Cas9 genome editing (Sup. Fig. 2B&4A). Lenalidomide and pomalidomide-induced degradation of FAM83F evident in DLD-1 wild-type cells was completely abolished in DLD-1 CRBN^-/-^ cells (Fig. 4B&C). Restoration of DLD-1 CRBN^-/-^ cells with human FLAG-CRBN partially restored the lenalidomide and pomalidomide-induced degradation of FAM83F. However, when DLD-1 CRBN^-/-^ cells were rescued with the FLAG-CRBN^V388I^ mutant, which mimics the mouse variant shown to be inactive for IMiD-induced protein degradation as the IMiD is unable to bind CRBN^V388I^ (7), lenalidomide and pomalidomide did not induce FAM83F degradation (Fig. 4B&C). These findings confirm that IMiD-induced FAM83F degradation requires the Cul4A^CRBN^ E3 ligase activity and is mediated by the proteasome.

**Figure 4:**
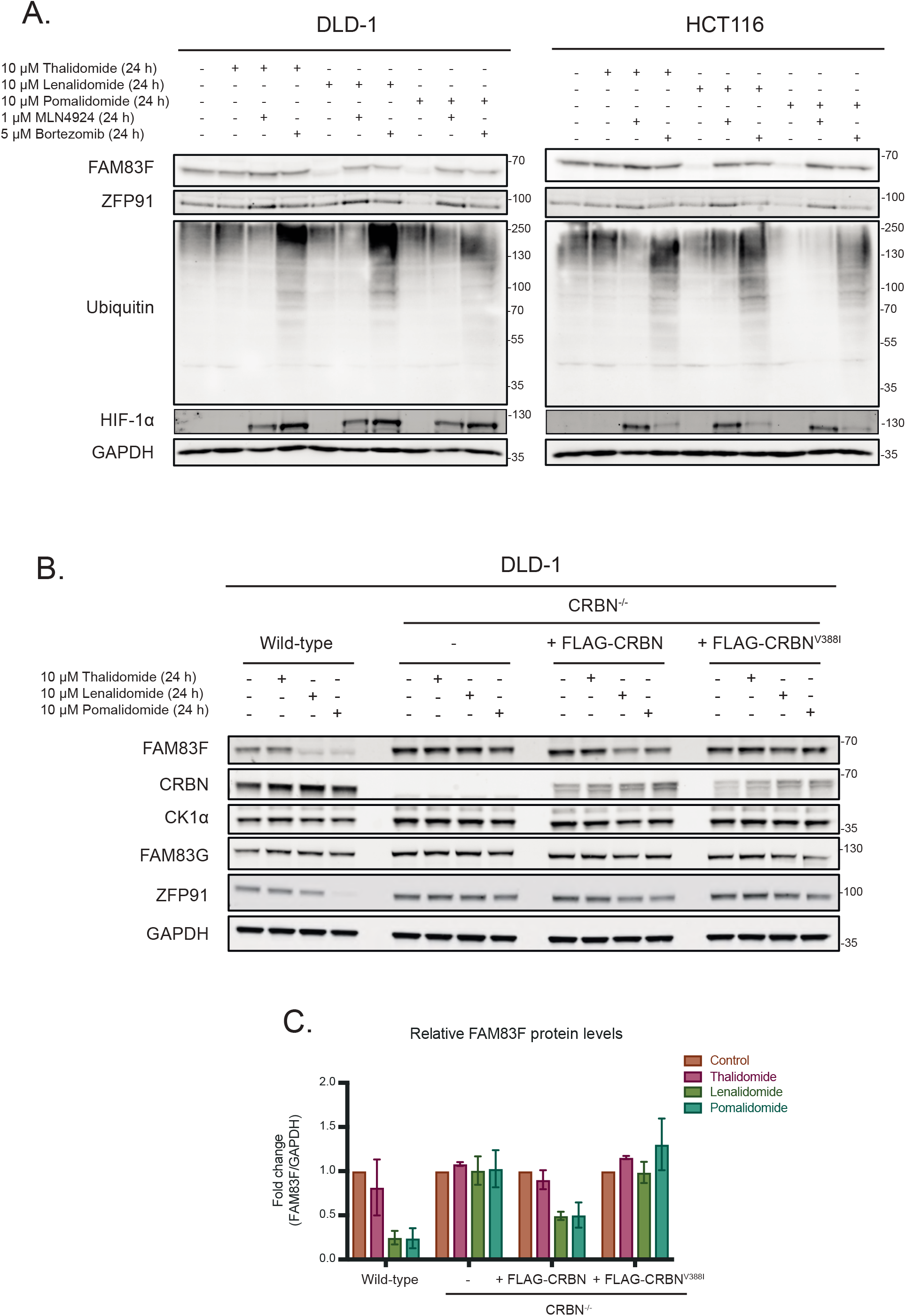
IMiD-induced FAM83F degradation occurs via the proteasome and is dependent on cereblon. **(A)** DLD-1 and HCT116 cell extracts treated with 10 μM IMiD compounds, 1 μM MLN4924, 5 μM Bortezomib or a combination thereof as indicated for 24 h were resolved by SDS-PAGE and subjected to Western blotting with the indicated antibodies. The accumulation of HIF-1α and ubiquitylated proteins following MLN4924 and Bortezomib treatments respectively were used as positive controls for successful compound treatments. **(B)** DLD-1 wild-type, DLD-1 CRBN^-/-^, DLD-1 CRBN^-/-^ rescued with FLAG-CRBN and DLD-1 CRBN^-/-^ rescued with FLAG-CRBN^V388I^ cell extracts treated with IMiD compounds (10 μM 24 h), were resolved by SDS-PAGE and subjected to Western blotting with the indicated antibodies. **(C)** Densitometry of FAM83F protein abundance upon treatment with IMiD compounds (10 μM 24 h) from (B). FAM83F protein abundance was normalised to GAPDH protein abundance and represented as fold change compared to untreated cells. Data representative of two biological replicates with bar graph representing mean ± standard error.

### FAM83G protects CK1α from IMiD-induced degradation

Lenalidomide-induced degradation of CK1α has been shown to be robust in multiple myeloid cells (7). In agreement, we observed more robust degradation of CK1α upon lenalidomide and other IMiD treatments in MV4.11 cells compared to DLD-1 cells (Sup. Fig. 5) or a panel of other cancer cell lines (Fig. 1). Interestingly, the levels of most FAM83 proteins in MV4.11 cells were either absent (FAM83B & F), or much lower in abundance (FAM83D, G & H) compared to DLD-1 cells (Sup. Fig. 5). We hypothesised that the absence of specific FAM83-CK1α complexes may explain why CK1α degradation with lenalidomide was more robust in MV4.11 cells compared to non-hematopoietic cell lines. To investigate whether specific FAM83 proteins impact IMiD-induced CK1α degradation, we generated HCT116 FAM83F^-/-^ and FAM83G^-/-^ cells with CRISPR/Cas9 genome editing (Sup. Fig. 2-4). Both lenalidomide and pomalidomide treatment caused FAM83F degradation in HCT116 wild-type and FAM83G^-/-^ cells, while FAM83G levels remained unchanged upon IMiD treatment in both HCT116 wild-type and FAM83F^-/-^ cells (Fig. 5A&B). Lenalidomide reduced CK1α protein abundance in HCT116 wild-type, FAM83F^-/-^ and FAM83G^-/-^ cell lines, but interestingly the reduction in FAM83G^-/-^ cells was slightly but significantly higher than wildtype cells (Fig. 5A&B). Additionally, thalidomide and pomalidomide were also able to induce a more robust CK1α degradation in FAM83G^-/-^ cells compared to wild-type and FAM83F^-/-^ cells (Fig. 5A&B). Considering this increased degradation is observed with multiple IMiDs in FAM83G^-/-^ cells, the possibility that, in the absence of FAM83G, an increased proportion of CK1α binds to FAM83F and is co-degraded as a complex was tested. Indeed, when we overexpressed wild-type GFP-FAM83G in HCT116 cells, pomalidomide-induced degradation of FAM83F was partially rescued suggesting that less CK1α is bound to FAM83F with CK1α preferentially binding to GFP-FAM83G (Fig. 5C-F). Thus, FAM83F is partially protected from degradation. Indeed, overexpression of GFP-FAM83G^F296A^, which has minimal interaction with CK1α (Fig 5C&D) (14) in HCT116 cells was unable to reduce the pomalidomide-induced degradation of FAM83F (Fig. 5E&F). Intriguingly, FAM83G was identified by mass spectrometry as one of the top interactors of CRBN upon IMiD treatment (2). This would suggest that multiple CK1α-FAM83 complexes could be recruited to the Cul4A^CRBN^ complex after exposure to IMiDs, but only the CK1α-FAM83F complex is positioned in such a way that ROC1 can ubiquitylate the both CK1α and FAM83F.

**Figure 5:**
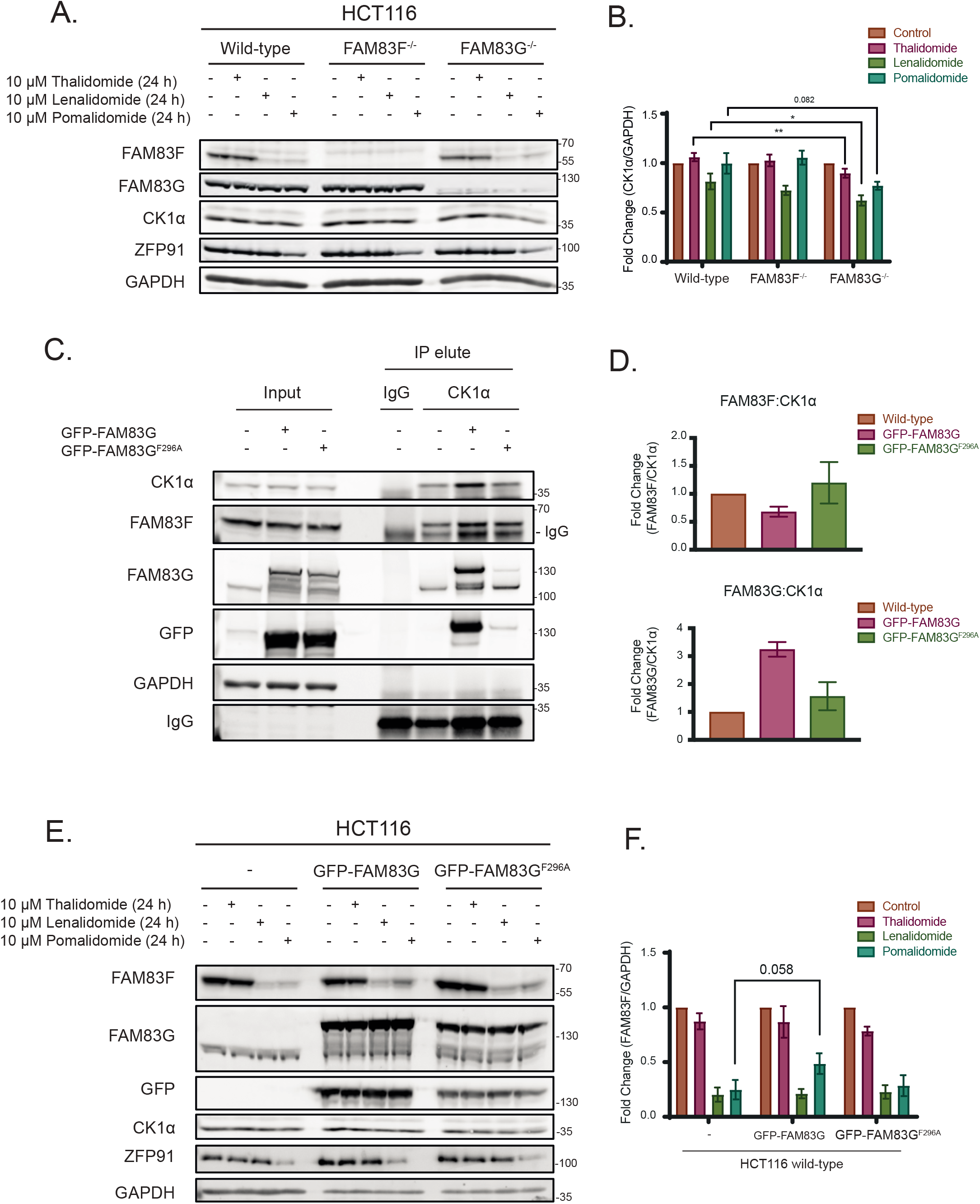
FAM83G protects CK1α from IMiD-induced degradation. **(A)** HCT116 wild-type, HCT116 FAM83F^-/-^ and HCT116 FAM83G^7^’ cell extracts treated with IMiD compounds (10 μM 24 h) were resolved by SDS-PAGE and subjected to Western blotting with indicated antibodies. **(B)** Densitometry of CK1α protein abundance upon treatment with IMiD compounds in HCT116 wild-type, HCT116 FAM83F^-/-^ and HCT116 FAM83G^-/-^ cells from (A). CK1α protein abundance was normalised to GAPDH protein abundance and represented as fold change compared to untreated cells. Data representative of seven biological replicates with bar graph representing mean ± standard error. **(C)** HCT116 wild-type, HCT116 wild-type transfected with GFP-FAM83G and HCT116 wild-type transfected with GFP-FAM83G^F296A^ cell extracts were subjected to IP with anti-CK1α antibody. Input extracts, and indicated IP elutes were resolved by SDS-PAGE and subjected to Western blotting with the indicated antibodies. **(D)** Densitometry of FAM83F and FAM83G protein abundance in anti-CK1α IP elute from (C). FAM83F and FAM83G protein abundances were normalised to CK1α protein abundance and represented as fold change compared to un-transfected HCT116 wild-type cells. Data representative of two biological repeats with bar graph representing mean ± standard error. **(E)** HCT116 wild-type, HCT116 wild-type transfected with GFP-FAM83G and HCT116 wild-type transfected with GFP-FAM83G^F296A^ cell extracts treated with IMiD compounds (10 μM 24 h) were resolved by SDS-PAGE and subjected to Western blotting with the indicated antibodies. **(F)** Densitometry of FAM83F protein abundance upon treatment with IMiD compounds from (E). FAM83F protein abundance was normalised to GAPDH protein abundance and represented as fold change compared to untreated cells. Data are representative of four biological replicates with bar graph representing mean ± standard error. Statistical analysis of (B) and (F) data was completed using a students unpaired t-test and comparing fold-change between untreated and IMiD treated samples. Statistically significant p-values are denoted as asterisks (**** <0.0001, *** <0.001, ** <0.01, * <0.05).

### BTX161 is an efficient CK1α-degrader which reduces FAM83G protein abundance through FAM83G-CK1α co-stability

The success of lenalidomide and pomalidomide in multiple myeloma treatment has led to the continuing development of new IMiD derivatives. BTX161 has been reported to be the most efficient CK1α degrader to date (23). We compared BTX161 to other IMiD compounds and confirmed the potency of BTX161 in CK1α degradation in MV4.11 and DLD-1 cells (Sup. Fig. 5). We sought to establish the effect of BTX161 on the FAM83 proteins using HCT116, DLD-1 and MV4.11 cells (Sup. Fig. 5 and Fig. 6A&B). We observed BTX161-induced degradation of FAM83F in DLD-1 and HCT116 cells (Sup. Fig. 5 and Fig. 6A&B). FAM83F and CK1α degradation in these cells occurred in a time and dose-dependent manner with optimal degradation observed after treatment with 10 μM BTX161 for 24 h (Fig. 6A&B). CK1α degradation was even more substantial in MV4.11 cells, which do not contain detectable levels of FAM83F protein (Sup. Fig. 5). We quantified protein levels of CK1α, FAM83F, FAM83H, FAM83G and FAM83B via Western blot analysis after treatment of HCT116, DLD-1 and MV4.11 cells with 10 μM BTX161 for 24 h (Fig. 6C). FAM83F and CK1α abundances were consistently reduced in all cell lines. FAM83H protein abundance was slightly reduced in HCT116 and MV4.11 cells. FAM83B protein abundance was unaffected by BTX161 treatment in all cells. FAM83G protein abundance was slightly reduced in DLD-1 cells but almost fully depleted in MV4.11 cells, while protein levels in HCT116 cells appear unchanged (Fig. 6C). We confirmed that this strong FAM83G degradation observed in MV4.11 cells was specific for BTX161 and, to a lesser extent, lenalidomide treatments (Sup. Fig. 6).

**Figure 6:**
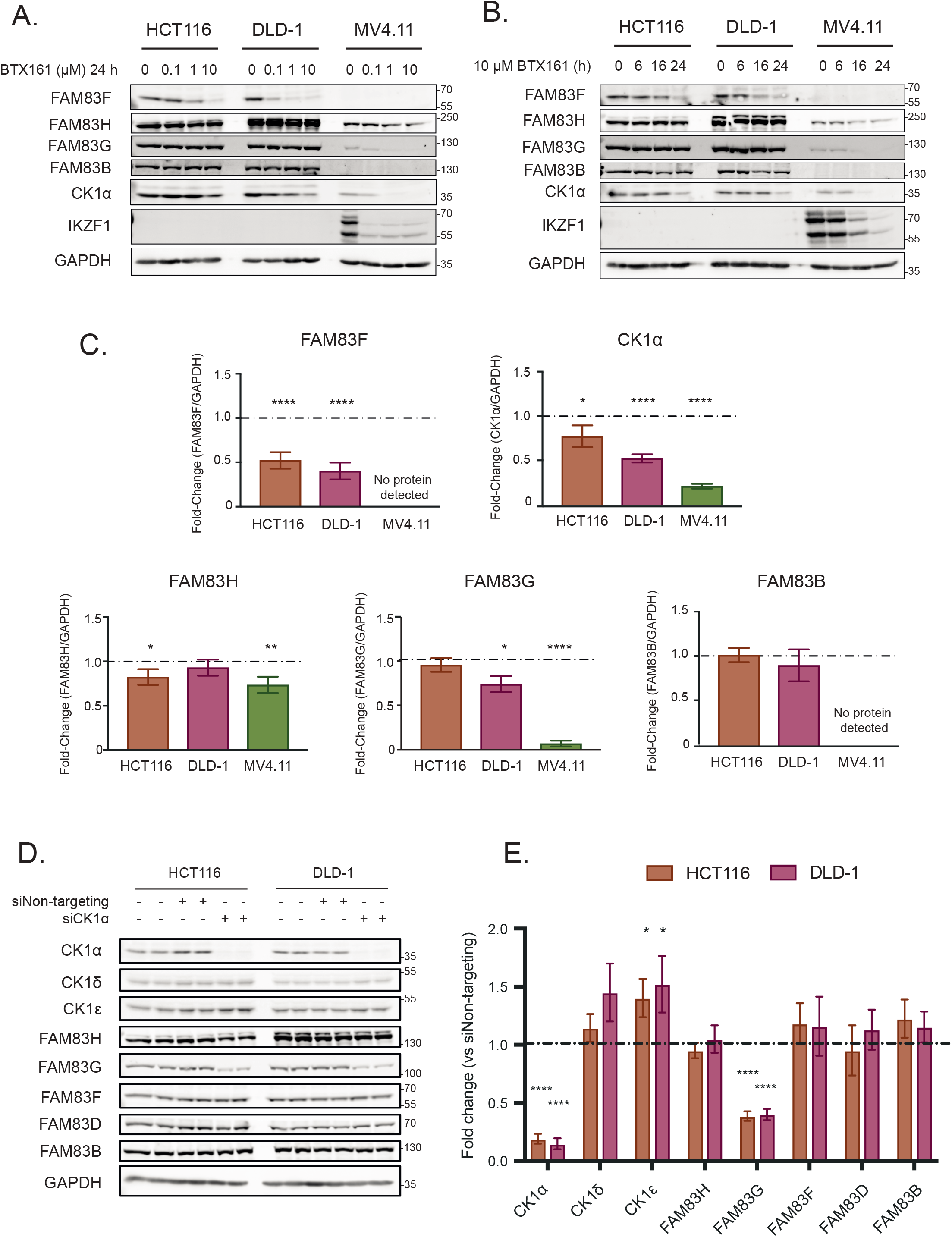
BTX161 is an efficient CK1α-degrader which reduces FAM83G protein abundance through FAM83G-CK1α co-stability. **(A)** HCT116, DLD-1 and MV4.11 cell extracts treated with varying concentrations of BTX161 (0, 0.1, 1 or 10 μM) for 24 h were resolved by SDS-PAGE and subjected to Western blotting with the indicated antibodies. **(B)** As in (A) except cells were treated with 10 μM BTX161 for varying times (0, 6, 16 or 24 h) prior to lysis. **(C)** Densitometry of FAM83H, FAM83G, FAM83F, FAM83B and CK1α protein abundance from (A & B) normalised to GAPDH protein abundance and represented as fold change compared to untreated cells. Data are representative of six biological replicates with bar graph representing mean ± standard error. The dashed line indicates a fold-change of 1 which is equivalent to an untreated sample. Statistical analysis was completed using a students unpaired t-test and comparing fold-change between untreated and samples treated with 10 μM BTX161 for 24 h. **(D)** HCT116 and DLD1 cell extracts transfected with control siRNA or siCK1α for 48 h were resolved by SDS-PAGE and subjected to Western blotting with the indicated antibodies. **(E)** Densitometry of CK1α, CK1δ, CK1ε, FAM83H, FAM83G, FAM83F, FAM83D and FAM83B protein abundance from (D) normalised to GAPDH protein abundance and represented as fold change compared to HCT116 and DLD-1 cells transfected with Non-targeting siRNA control. Data representative of three biological repeats with bar graph representing mean ± standard error. The dashed line indicates a fold-change of one which is equivalent to an untreated sample. Statistical analysis was completed using a students unpaired t-test and by comparing fold-change between cells transfected with Non-targeting siRNA control and cells transfected with siCK1α. Statistically significant p-values are denoted as asterisks (**** <0.0001, *** <0.001, ** <0.01, * <0.05).

Given that the reductions in FAM83G protein abundance in various cell lines mirror the reduction in CK1α protein, we queried whether the FAM83G-CK1α complex is co-degraded by BTX161, similarly to the FAM83F-CK1α complex, or if this was a result of CK1α-dependent stability of FAM83G. CK1α protein was reduced in HCT116 and DLD-1 cells using CK1α-targeting small interfering RNA (siCK1α) (Fig. 6D&E). This produced an approximate 75-80% reduction in CK1α protein after 48 h and resulted in a significant reduction in FAM83G protein when compared to non-targeting siRNA controls (Fig. 6D&E). Loss of CK1α, caused by either BTX161 treatment or siRNA, resulted in reduced FAM83G protein levels suggesting some form of co-stability between the two proteins. In contrast, FAM83F protein levels were unaffected by siRNA treatment. We therefore propose that only FAM83F-CK1α complex can be degraded by lenalidomide, pomalidomide and BTX161 but the loss of CK1α by lenalidomide and BTX161 can result in a reduction in FAM83G due to co-stability. The efficiency of CK1α degradation by IMiDs varies between cell lines and may be affected by the abundance of FAM83 proteins, some of which may protect CK1α from IMiD-induced degradation and thus the abundance of specific FAM83 proteins in cells could predict the efficacy of overall CK1α degradation.

### Inducible degradation of FAM83F attenuates Wnt signalling

We have recently established that FAM83F mediates canonical Wnt signalling through association with CK1α (20). Ablation of FAM83F significantly inhibits Wnt signalling in multiple cell lines. The plasma membrane localisation of FAM83F mediated by its C-terminal farnesylation is also essential for its role in Wnt signalling (20). Therefore, we sought to investigate the role of IMiD-induced FAM83F degradation on canonical Wnt signalling, by assessing the abundance of a Wnt-target gene, *Axin2* (24), in DLD-1 wild-type, FAM83F^-/-^ and CRBN^-/-^ cells following pomalidomide treatment (Fig. 7A). We chose DLD-1 cells because they harbour a truncated adenomatous polyposis coli (APC) protein mutant and display hyperactive Wnt signalling (25). Additionally, IMiD treatment of DLD-1 cells efficiently reduced FAM83F, but not CK1α protein levels (Fig. 4B). Treatment of DLD-1 cells with 10 μM pomalidomide for 48 h significantly reduces *Axin2* mRNA abundance in wild-type cells. Pomalidomide treatment does not alter *Axin2* mRNA abundance in DLD-1 FAM83F^-/-^ or DLD-1 CRBN^-/-^ cells indicating that the pomalidomide-induced reduction in *Axin2* mRNA requires FAM83F and CRBN. To confirm that treatment with pomalidomide could replicate all phenotypes associated with genetic knockout of FAM83F, membrane fractions from DLD-1 wild-type, FAM83F^-/-^ and CRBN^-/-^ cell lines treated with 10 μM pomalidomide for 24 h were assessed for FAM83F and CK1α protein abundance (Fig. 7B). In DLD-1 wild-type cells, membranous FAM83F and CK1α protein abundance was reduced upon pomalidomide treatments whilst no changes were detected in DLD-1 CRBN^-/-^ cells. CK1α levels in membrane fractions of DLD-1 FAM83F^-/-^ cells, which were already reduced under untreated conditions compared to DLD-1 wild-type cells, were not altered upon pomalidomide treatment. These results demonstrate that IMiD-induced degradation of FAM83F protein replicate the phenotypes observed with genetic FAM83F knockout cells and, importantly, IMiD-induced degradation of FAM83F appears to reduce Wnt activity in colorectal cancer cells displaying constitutively active Wnt signalling.

**Figure 7:**
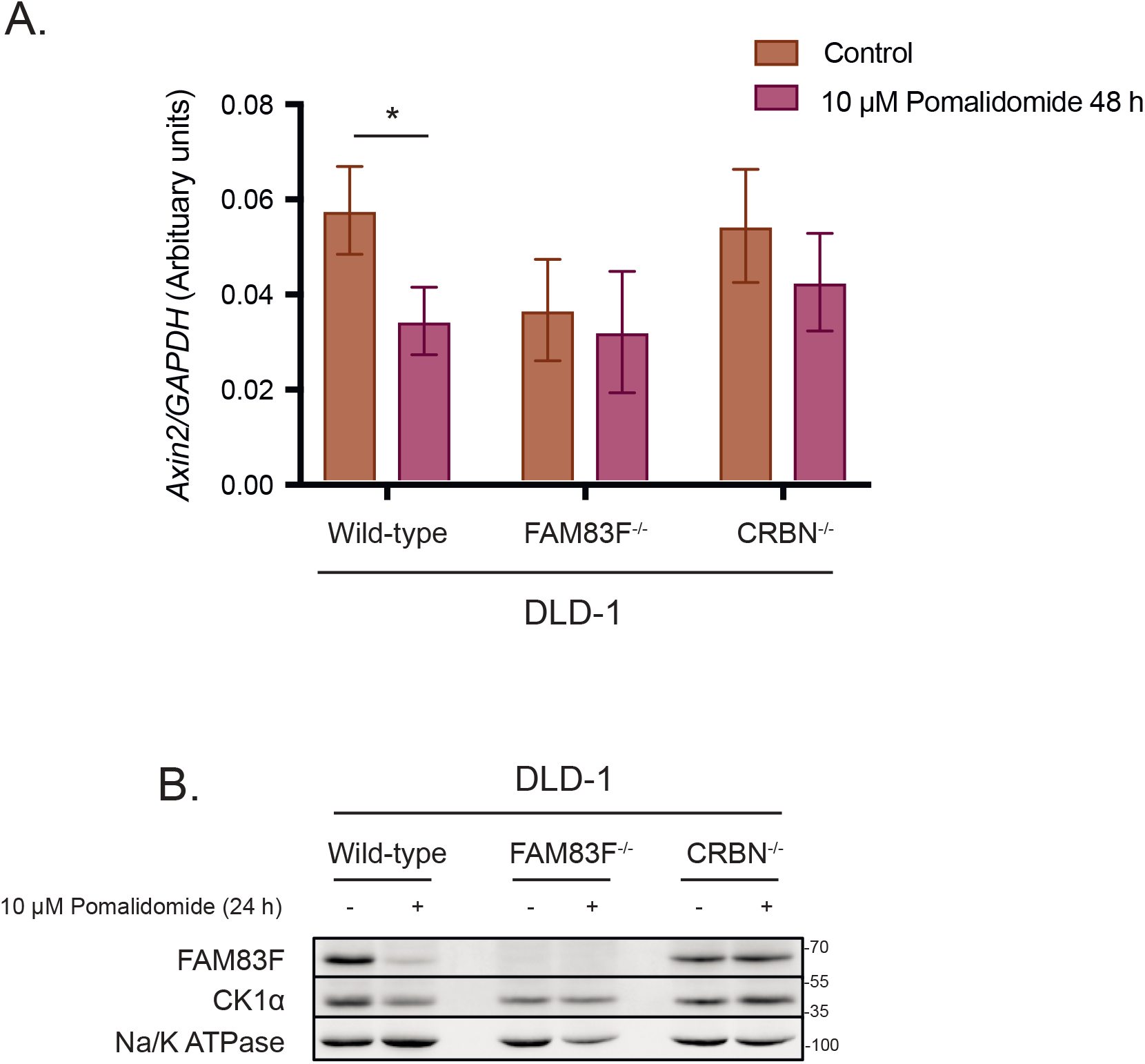
IMiD-induced degradation of FAM83F attenuates Wnt signalling and removes CK1α from the plasma membrane. **(A)** qRT-PCR was performed using cDNA from DLD-1 wild-type, DLD-1 FAM83F^-/-^ and CRBN^-/-^ cell lines following treatment with 10 μM pomalidomide for 48 h, and primers for *Axin2* and *GAPDH* genes. *Axin2* mRNA expression was normalised to *GAPDH* mRNA expression and represented as arbitrary units. Data are representative of five biological replicates with bar graph representing mean ± standard error. Statistical analysis was completed using a students unpaired t-test and by comparing cell lines as denoted on graph. p-values denoted by asterisks (**** <0.0001, *** <0.001, ** <0.01, * <0.05). **(B)** Membrane fractions from DLD-1 wild-type, DLD-1 FAM83F^-/-^ and DLD-1 CRBN^-/-^ cell lines, following treatment or not with 10 μM pomalidomide for 24 h, were separated by SDS-PAGE and subjected to Western blotting with the indicated antibodies. The specificity of membrane compartment isolation was determined with Western blotting for Na/K ATPase, a membrane specific protein.

## DISCUSSION

Lenalidomide (Revlimid) was the second highest grossing prescription drug worldwide in 2018 and with clinical trials for additional haematological derived cancers ongoing, the prominence of IMiDs are likely to increase further (26). Therefore, the discovery of novel proteins targeted for IMiD-induced degradation is important to predict both unforeseen consequences of IMiD treatments and potential new therapeutic targets. Here, we report that several IMiD compounds can induce degradation of the FAM83F protein and demonstrate that degradation of FAM83F requires its ability to interact with CK1α, indicating that IMiD-induced recruitment of CK1α to CRBN mediates the co-recruitment of FAM83F (Fig. 8). It remains unknown whether all FAM83-CK1α complexes are recruited to the Cul4A^CRBN^ complex upon IMiD treatment but not all complexes are ubiquitylated and degraded. Alternatively, the binding of certain FAM83 proteins to CK1α could restrict the recruitment of CK1α to CRBN. However, given that FAM83G was reported as one of the top binders of CRBN from cells treated with lenalidomide (2), the former hypothesis appears more likely. Mass spectrometry has identified that the abundance of other FAM83 proteins can be reduced upon IMiD treatment (27) which we have not observed with our treatments with the exception of FAM83G after BTX161 treatment. However, given the substantial CK1α degradation observed upon BTX161 treatment and that knockdown of CK1α through siRNA reduces FAM83G abundance, these observations on FAM83G protein stability are likely a result of FAM83G-CK1α co-stability rather than direct complex degradation.

**Figure 8:**
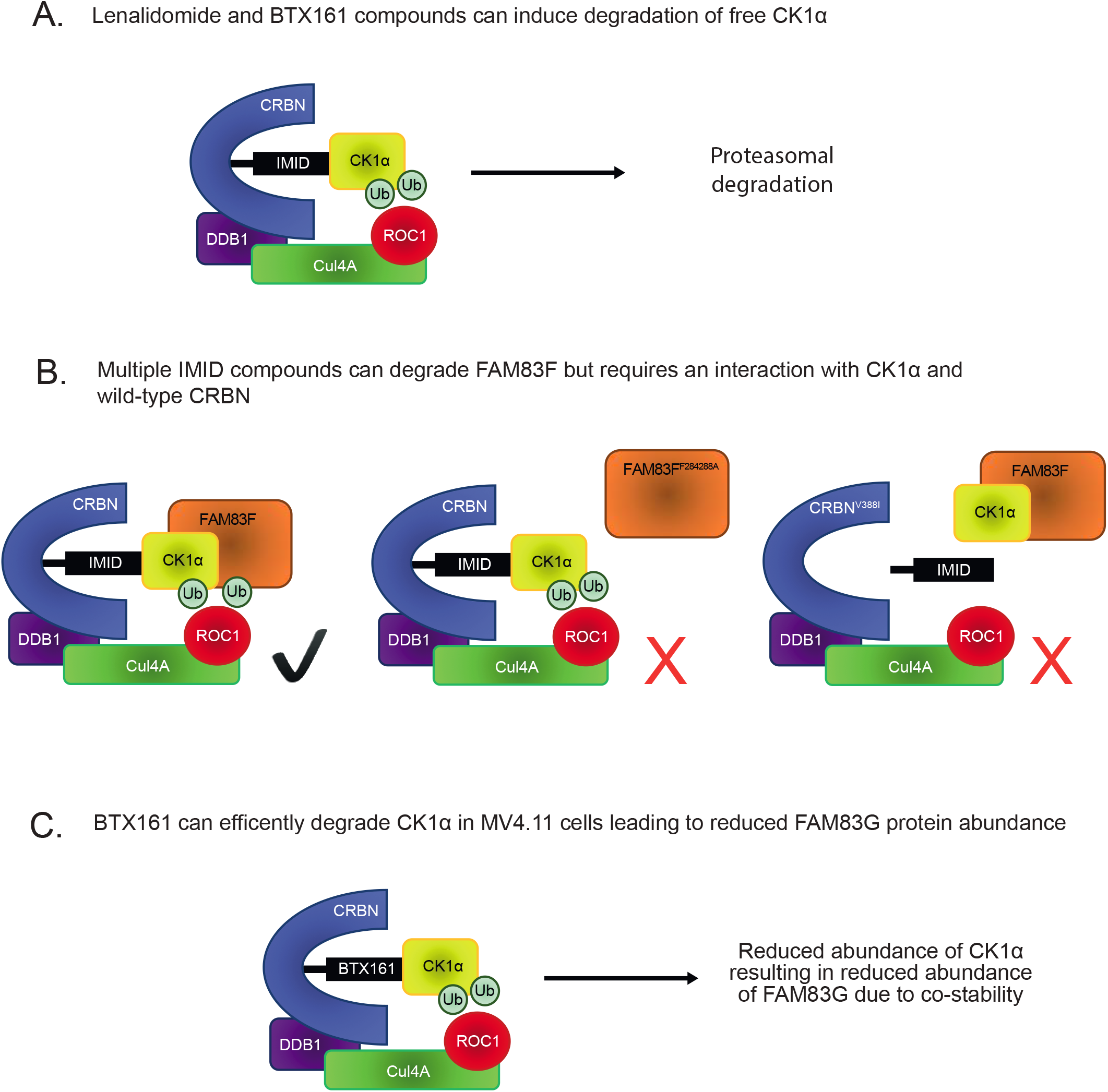
Proposed model for IMiD-induced FAM83F and CK1α degradation. **(A)** Previously reported mode of action for CK1α degradation by lenalidomide and BTX161 compounds. The IMiD can bind CK1α and a conserved binding pocket in CRBN, thus bringing CK1α into proximity of the Cul4A^CRBN^ complex, facilitating the addition of ubiquitin by ROC1 to CK1α. CK1α is then subsequently degraded via the proteasome. **(B)** Our proposed model for FAM83F degradation by multiple IMiD compounds. The IMiDs can bind the CK1α-FAM83F complex and a conserved binding pocket in CRBN, thus bringing the CK1α-FAM83F complex into proximity of the Cul4A^CRBN^ complex, facilitating the addition of ubiquitin by ROC1 to the CK1α-FAM83F complex. CK1α and FAM83F are then subsequently degraded via the proteasome. Mutation of FAM83F at F284 and F288 to alanines abolishes the interaction with CK1α, thus there is no recruitment or subsequent degradation of FAM83F upon IMiD treatment. Mutation of CRBN at V388 to isoleucine to mimic the mouse homolog which cannot bind IMiDs prevents IMiD from binding to CRBN, thus no IMiD neo-substrate, including the CK1α-FAM83F complex, can be recruited to the Cul4A^CRBN^ complex. **(C)** We propose that the BTX161-induced reduction in FAM83G protein abundance is a result of efficient CK1α degradation by BTX161 resulting in a loss of FAM83G protein due to FAM83G-CK1α co-stability, and not direct BTX161-induced FAM83G degradation.

Increasingly, targeted protein degradation is being used as an investigative tool in cell biology, with hopes to translate into a therapeutic option for a variety of diseases (28). The efficacy of inducible protein degradation is influenced by a number of factors including protein synthesis rate, binding affinity of protein of interest to E3 ligase complexes, efficiency of ubiquitylation and presence of deubiquitinases (29). Ubiquitylation requires accessible lysine residues in close proximity to the E3 ligase (30). IMiDs bind neo-substrates at slightly different angles, thus presenting different residues to the ROC1 E3 ligase which may explain the differences in protein degradation noted between various IMiD compounds (31). However, numerous unresolved questions remain regarding targeted proteolysis, including how the formation of different protein complexes affects degradation kinetics and/or occurrence. With our data, we make important observations suggesting that the architecture of CK1α in different FAM83-CK1α complexes most likely determines whether, and to what extent, CK1α and associated FAM83 protein can be degraded by IMiDs. Specifically, the FAM83F-CK1α complex is robustly degraded by various IMiD compounds, whilst sequestering CK1α in the FAM83G-CK1α complex spares CK1α from IMiD-induced degradation. Clinically, lenalidomide treatment in del(5q) MDS patients was shown to be effective and it was suggested that this might be due to haploinsufficiency of CK1α (7). Our data suggests that the efficiency of CK1α degradation is rather influenced by the relative abundance of FAM83 proteins and suggests that FAM83 protein abundance may be used as predicting biomarkers of IMiD-induced CK1α degradation, which may widen the potential use of IMiDs. In addition to IMiDs, other target protein degraders such as PROTACs are emerging as key therapeutic modalities in drug research (32). Our observations presented in this study clearly illustrate that the nature of the inherent complex in which the target protein exists is an important factor that determines whether the target protein can be degraded. This not only provides challenges in trying to design degraders of specific target proteins that yield complete degradation but also provide opportunities in which specific protein complexes can be targeted for degradation thereby affecting specific functions of target proteins. In this regard, the efficacy for a protein degrader should perhaps not be judged by how much a target protein is degraded but rather to what extent a change in expected phenotype is achieved. Indeed, for proteins which exist in distinct functional pools, specific degradation of subcomplexes may be sufficient to disrupt target pathways, whilst leaving other non-targeted subcomplex functions intact.

FAM83F has been implicated in oncogenesis with high FAM83F expression observed in oesophageal squamous cell carcinoma (33), lung adenocarcinoma (34), glioma (35) and thyroid carcinoma (36). In contrast, FAM83F has been reported to enhance the stabilisation and activity of the tumour suppressor p53 with siRNA targeting FAM83F reducing cellular proliferation (37). These effects are dependent on the mutational status of p53 with overexpression of FAM83F increasing cell migration in cells containing mutant p53, indicating FAM83F may promote or inhibit cancer progression depending on the tumour’s mutational status. Recently, we have demonstrated that the membranous FAM83F-CK1α complex activates canonical Wnt signalling (20). Activated Wnt signalling is a hallmark of many cancers, especially CRCs (38). However, it is often believed that genetic alterations which activate canonical Wnt signalling in sporadic CRCs, specifically those at the level of the β-catenin destruction complex, will render any inhibition of upstream membranous Wnt signalling proteins ineffectual (39). In contrast to this, we demonstrate that reducing membrane-associated FAM83F-CK1α with IMiD treatment can modulate canonical Wnt signalling in DLD-1 cells, which contain mutant APC and are unresponsive to Wnt3A stimulation. Lenalidomide has been tested in phase I and phase II clinical trials for sporadic CRCs but clinical response has been poor (40). Often these trials are in advanced metastatic tumours which contain multiple oncogenic driver mutations and aberrant signalling pathways. The clinical effect of IMiDs on an early Wnt-dependent disease such as the initial polyp formation in Familial Adenomatous Polyposis (FAP), which is caused by mutant Apc (41), may yield more promising results.

## MATERIALS & METHODS

### Plasmids

All constructs are available for request from the MRC-PPU reagents website (http://mrcppureagents.dundee.ac.uk). The unique identifier (DU) numbers provide direct links to the cloning strategy and sequence information. Sequences were verified by the DNA sequencing service, University of Dundee (http://www.dnaseq.co.uk). Constructs generated include: pBABED.puro FAM83F (DU37979), pBABED.puro FAM83F^F284A/F288A^ (DU28196), pBABED.puro FLAG CRBN (DU54685), pBABED.puro FLAG CRBN^V388I^ (DU64137), pBABED.puro U6 FAM83F tv1 Nter KI sense (DU54050), pX335 FAM83F Nter KI Antisense (DU54056), pMS-RQ FAM83F Nter GFP donor (DU54325), pBABED.puro U6 FAM83F ex2 KO sense (DU54848), pX335 FAM83F ex2 KO Antisense (DU54850), pBABED.puro U6 CRBN ex3 KO sense A (DU64046), pX335 CRBN ex3 KO Antisense A (DU64483), pBABED.puro U6 FAM83G ex2 KO sense (DU52480), pX335 FAM83G ex2 KO Antisense (DU52484), pcDNA5-FRT/TO-GFP-FAM83G (DU33272) and pcDNA5-FRT/TO-GFP-FAM83G^F296A^ (DU28477).

Plasmid amplification was completed by transforming 10 μl *E. coli* DH5α competent cells (Invitrogen) using 1 μl of plasmid DNA. Bacteria were incubated on ice for 10 min before heat-shocking at 42°C for 45 s. Following a further 2 min on ice, the transformed bacteria were plated on LB-agar medium plates containing 100 μg/ml ampicillin and incubated at 37°C for 16 h. Single colonies were used to inoculate a 5 ml culture of LB medium containing 100 μg/ml ampicillin then incubated at 37°C for 16 h with constant shaking. Plasmid DNA was purified from a bacterial culture using Qiagen mini-prep kit by following the manufacturer’s protocol. Isolated DNA yield was quantified using a Nanodrop 1000 spectrophotometer (Thermo Fisher Scientific).

### Antibodies

Antibodies recognising FAM83B (SA270), FAM83D (SA102), FAM83F (SA103), FAM83H (SA273), CK1α (SA527), CK1ε (SA610), CK1δ (SA609) and GFP (S268B) were generated in-house and are available for request from the MRC-PPU reagents website (http://mrcppureagents.dundee.ac.uk). Antibodies recognising GAPDH (14C10) (#2118), IKZF1 (D6N9Y) (#14859), CRBN (D8H3S) (#71810), Na, K-ATPase alpha1 (D4Y7E) (#23565) and Lamin A/C (#2032) were obtained from Cell signalling technology. Additional antibodies used were FAM83G (ab121750, Abcam), Alpha-tubulin (MA1-80189, Thermo Fisher Scientific), Ubiquitin (BML-PW8810, Enzo), HIF-1α (6109590, BD biosciences) and ZFP91 (A303-245A, Bethyl laboratories). Secondary antibodies used were StarBright Blue 700 goat anti-rabbit IgG (12004161, BioRad), StarBright Blue 700 goat anti-mouse IgG (12004158, BioRad), IRDye 800CW donkey anti-goat IgG (926-32214, Licor) and IRDye 800CW goat anti-rat IgG (926-32219, Licor).

### Cell culture

THP-1 (TIB-202, ATCC) and MV4.11 (CRL-9591, ATCC) cells were maintained in Roswell Park Memorial Institute 1640 medium (RPMI; Gibco). HCT116 (CCL-247, ATCC), DLD-1 (CCL-221, ATCC), PC-3 (CRL-1435, ATCC), A549 (CCL-185, ATCC), HaCaT (from Joan Massague’s lab at Memorial Sloan Kettering Cancer Centre, not commercially obtained but can be provided on request) (42) and HEK293-FT (R70007, Thermo Fisher Scientific) cells were maintained in Dulbecco’s Modified Eagle’s Medium (DMEM; Gibco). RPMI and DMEM were supplemented with 10% (v/v) FBS (F7524, Sigma), 2 mM L-glutamine (25030024, Invitrogen) 100 units/ml penicillin and 100 mg/ml streptomycin (15140122, Invitrogen). Cells lines were regularly tested for mycoplasma contamination and only mycoplasma free cell lines were used for experimentation.

### Generation of ^GFP/GFP^FAM83F, FAM83F^-/-^, CRBN^-/-^ and FAM83G^-/-^ cell lines using CRISPR/Cas9 genome editing

For the generation of FAM83F knock-out HCT116 and DLD-1 cell lines, the *FAM83F* locus was targeted with a dual guide RNA approach using the sense guide RNA (pBabeD-puro vector, DU54848); GCGTCCAGGATGATGTACACT and antisense guide RNA (pX335-Cas9-D10A vector, DU54850); GGCAGGAGTGAAGTATTTCC. For the generation of CRBN knock-out DLD-1 cell lines, the *CRBN* locus was targeted with a dual guide RNA approach using the sense guide RNA (pBabeD-puro vector, DU64046); GCTCAAGAAGTCAGTATGGTG and antisense guide RNA (pX335-Cas9-D10A vector, DU64483); GTGAAGAGGTAATGTCTGTCC. For the generation of FAM83G knock-out HCT116 cell lines, the *FAM83G* locus was targeted with a dual guide RNA approach using the sense guide RNA (pBabeD-puro vector, DU52480); GGACCGCTCCATCCCGCAGC and antisense guide RNA (pX335-Cas9-D10A vector, DU52484); GCTGGGGCCAGTACTCCAGGG. For generation of N-terminal GFP knock-in to the *FAM83F* locus, the *FAM83F* locus was targeted with a dual guide RNA approach using the sense guide RNA (pBabeD-puro vector, DU54050); GTTCAGCTGGGACTCGGCCA, antisense guide RNA (pX335-Cas9-D10A vector, DU54056); GCGAGGCGCACGTGAACGAGA and the GFP-FAM83F donor (pMK-RQ vector, DU54325).

Plasmids (1 μg of sense and antisense guide RNAs + 3 μg donor for knockins) were diluted in 1 ml OptiMem (Gibco) and 20 μl of polyethylenimine (PEI; 1 mg/ml) (Polysciences) was added. This transfection mix was vortexed vigorously for 15 s, incubated for 20 min at room temperature and then added dropwise to a 10 cm diameter dish containing approximately 70% confluent cells in complete culture medium. Selection of transfected cells was performed 24 h post transfection in medium containing 2 μg/ml puromycin for 48 h. Single cells were isolated by fluorescence-activated cell sorting (FACS), with single GFP-positive cells (for knock-ins) or all cells (for knock-outs) isolated, and plated into individual wells of 96-well plates, pre-coated with 1% (w/v) gelatin (Sigma). Viable clones were expanded and assessed for successful knock-in or knock-out by both Western blotting and genomic DNA sequencing (Sup. Fig. 2-4).

For verification by DNA sequencing, the region surrounding the gRNA target sites were amplified by PCR with KOD Hot Start Polymerase (Merck) according to manufacturer’s instructions with the following primer pairs: FAM83F exon 2 (Forward: TCATTGCTGTGGTCATGGAC, Reverse: AATCCGGAAGTCAGTGAGCT), FAM83F N-terminal GFP KI (Forward: TGTACAAGGCCGAGAGTCAGCTGAACTG, Reverse: CAGTTCAGCTGACTCTCGGCCTTGTACA), CRBN exon 3 (Forward: GGTGCTGATATGGAAGAATTTCATGGC, Reverse: GTATGAAGGTGAAGAGCTGAGTTAGATGG) and FAM83G exon 2 (Forward: TCTTTCCCGCAGATTGCTCATGG, Reverse: TTCTTCTGGGGAACCAGAAACACC). PCR products of positive clones were cloned with the StrataClone PCR Cloning Kit (Agilent) into the supplied vector system, according to the manufacturer’s protocol. Sequencing of the edited loci in the positive clones was performed by the MRC-PPU DNA sequencing and services (http://mrcppureagents.dundee.ac.uk).

### Transient transfections

Transient transfections were performed in HCT116 cells with either pcDNA5-FRT/TO-GFP-FAM83G (DU33272) or pcDNA5-FRT/TO-GFP-FAM83G^F296A^ (DU28477). Plasmids (1 μg) were diluted in 1 ml OptiMem (Gibco) and 20 μl PEI (1mg/ml) was added. The transfection mix was incubated for 20 min at room temperature, then added dropwise to a 10 cm diameter dish of cells in complete culture medium. Fresh media was added 24 h post transfection and indicated treatments performed thereafter.

### Retroviral transductions

Retroviruses were produced using the following constructs; pBABED.puro FAM83F^WT^ (DU37979), pBABED.puro FAM83F^F284A/F288A^ (DU28196), pBABED.puro FLAG CRBN (DU54685) and pBABED.puro FLAG CRBN^V388I^ (DU64137). Retroviruses were produced by transfecting HEK293-FT cells as previously described (16). Briefly, 6 μg pBabe plasmid, 3.8 μg pCMV5-GAG/Pol (Clontech), 2.2 μg pCMV5-VSV-G (Clontech) were diluted in 600 μl OptiMem (Gibco) and 24 μl PEI (1 mg/ml) was added. The transfection mixture was incubated for 20 min at room temperature then added dropwise to a 10 cm diameter dish of cells in complete culture medium. Fresh media was added 24 h post transfection. Media containing retroviruses was collected after 24 h and passed through a 0.45 μm sterile syringe filter. For transduction of target cells, 1 ml of retroviral medium together with 8 μg/ml polybrene (Sigma) was added to a 10 cm diameter dish of cells containing 9 ml complete culture medium. Selection of transduced cells was performed 24 h post transduction with cells incubated in media containing 2 μg/ml puromycin for 48 h. Successful transduction was assessed by Western blotting.

### Compound treatments

IMiD compounds (Thalidomide, Lenalidomide, Pomalidomide and Iberdomide) were obtained from Caymen Chemicals. BTX161 compound was synthesised in-house and is available for request from the MRC-PPU reagents website (http://mrcppureagents.dundee.ac.uk). IMiD compounds were added to cell culture media at indicated concentrations (between 0.1 μM to 10 μM) for indicated duration. dTAG-13 (Sigma), a CRBN-binding PROTAC, was used at 1 μM for 24 h. MLN4924 (Sigma), an inhibitor of NEDD8-activating E1 enzyme, was used at 1 μM for 24 h. Bortezomib (Sigma), a proteasome inhibitor, was used at 5 μM for 24 h.

### Cell lysis, SDS-PAGE and Western blotting

Cells were washed and scraped in ice-cold PBS, and then pelleted. Cell pellets were lysed in lysis buffer (20 mM Tris-HCl (pH 7.5), 150 mM NaCl, 1 mM EDTA, 1 mM EGTA, 1% (v/v) Triton X-100, 2.5 mM sodium pyrophosphate, 1 mM beta-glycerophosphate, 1 mM Na3VO4, 1x complete EDTA-free protease inhibitor cocktail (Roche)). Lysates were clarified at 13000rpm for 20 min. Protein concentration was measured using Pierce Coomassie Bradford protein assay kit (Thermo Fisher Scientific). Protein concentrations were adjusted to 1-3 μg/μl in lysis buffer and NuPAGE 4x LDS sample buffer (NP0007) (Thermo Fisher Scientific) was added to lysates. Lysates (20-30 μg protein) were separated by sodium dodecyl sulphate-polyacrylamide electrophoresis (SDS-PAGE) and gels were transferred to nitrocellulose membranes. Following blocking in 5% (w/v) milk in TBS-T (50 mM Tris-HCL (pH 7.5), 150 mM NaCl, 0.1% (v/v) Tween 20) for 60 min, membranes were incubated in primary antibody (1:1000 dilution in blocking buffer) for 16 h at 4°C, washed 3×10 min in TBS-T, then incubated in secondary antibody (1:5000 dilution in blocking buffer) for 1 h at room temperature and washed 3×10 min in TBS-T. Fluorescence of secondary antibody was detected using the Chemidoc system (BioRad) and data analysed by Image lab software (BioRad). Densitometry of protein blots was completed using Image J software (https://imagej.net). The density of protein of interest bands were measured and normalised to those of loading control bands with fold change calculations and statistical analysis performed using Microsoft Excel software (www.microsoft.com). Graphical representations of data were prepared using Prism 8 (www.graphpad.com).

### Subcellular fractionation

Cells were washed and scraped in ice-cold PBS, and then pelleted. Cellular fractionation into cytoplasmic, nuclear and membrane lysates was completed using a subcellular protein fractionation kit (Thermo Fisher Scientific) following manufacturer’s instructions. Briefly, cells were lysed in sequential buffers to separate cellular compartments into cytoplasmic, membrane, nuclear and cytoskeletal fractions. Quantification of protein concentrations, sample preparation and SDS-PAGE were completed as previously described.

### Immunoprecipitation

Protein lysates were prepared, and protein concentration quantified as previously outlined. Anti-CK1α antibody (1 μg) was added to each lysate sample (1 mg protein) and incubated on a rotating wheel for 16 h at 4°C. Protein G sepharose beads (DSTT) pre-equilibrated in cell lysis buffer were added (20 μl of 50% beads:lysis buffer slurry) to each lysate sample and incubated on a rotating wheel for 1 h at 4°C. Beads were pelleted and supernatant removed and stored as flow-through. Beads were washed in cell lysis buffer three times. Elution was performed by the addition of 20-40 μl of NuPAGE 1x LDS sample buffer to the beads, followed by denaturing proteins at 95°C for 10 min. Input and eluted samples were analysed by SDS-PAGE as previously described.

### Immunofluorescence

Cells were plated on sterile glass coverslips. Cells were fixed in 4% (v/v) paraformaldehyde for 15 min, permeabilized in 0.2% (v/v) Triton X-100 for 10 min then blocked in 5% (w/v) bovine serum albumin for 60 min. Cells were incubated in primary antibody (1:100) diluted in 0.5% (w/v) bovine serum albumin for 16 h at 4°C. Following washing in PBS/0.1% Tween-20, cells were incubated in Alexa Fluor 594 secondary (1:500; Thermo Fisher Scientific) for 60 min at room temperature. Coverslips were washed in PBS/0.1% Tween-20 and then incubated with 1 μg/ml DAPI (Sigma) for 5 min at room temperature to visualise nuclei. Coverslips were mounted on glass slides in Vectashield mounting media (Vector Laboratories) and sealed with CoverGrip coverslip sealant (Biotium). Images were captured on a Deltavision microscope using 60x objectives. Images were prepared using Omero software (www.openmicroscopy.org).

### Transfection with small interfering RNA (siRNA)

ON-TARGETplus human *CSNK1A1* siRNA (L-003957-00-0005, Dharmacon) and ON-TARGETplus Non-targeting pool (D-001810-10-05, Dharmacon) were resuspended using 5x siRNA buffer (B-002000-UB-100, Dharmacon). Adherent cells were seeded in 6-well plates and grown to 70-80% confluence. siRNA was diluted in OptiMem (Gibco) to a final working concentration of 25 pmol siRNA per well of a 6-well plate. Lipofectamine RNAi-MAX transfection reagent (13778100, Thermo Fisher) was diluted in OptiMem to a final working volume of 7.5 μl Lipofectamine per well. After incubating both siRNA and Lipofectamine samples separately for 5 min at room temperature, they were mixed and incubated for 20 min at room temperature. The siRNA-Lipofectamine mixture was added dropwise to cells and incubated for 48 h before lysis.

### Quantitative real time polymerase chain reaction (qRT-PCR)

Cells were grown to approximately 70% confluence in 6-well plates. Cells were then treated with IMiD compounds as indicated before RNA extractions were performed using the RNeasy mini kit (Qiagen). RNA was quantified using a NanoDrop 3300 Fluorospectrometer (Thermo Fisher Scientific). Synthesis of cDNA was completed using iScript cDNA synthesis kit (BioRad) and 1 μg of RNA per reaction. qRT-PCR was performed in triplicate in a 10 μl final volume with 2 μM forward primer, 2 μM reverse primer, 50% (v/v) iQ SYBR green supermix (BioRad) and 2 μl cDNA (diluted 1:5) using a CFX384 real-time system qRT-PCR machine (BioRad). Primers were designed using Benchling and purchased from Invitrogen. *Axin2* forward: TACACTCCTTATTGGGCGATCA, *Axin2* reverse: TTGGCTACTCGTAAAGTTTTGGT, *GAPDH* forward: TGCACCACCAACTGCTTAGC, *GAPDH* reverse: GGCATGGACTGTGGTCATGAG. The datasets were analysed using the comparative Ct method (ΔΔCt Method) (43) with *Axin2* the Wnt-target gene and *GAPDH* the endogenous control gene. Statistical analysis was performed using Microsoft Excel software (www.microsoft.com) and graphical representations of data were prepared using Prism 8 (www.graphpad.com).

## Acknowledgements

We thank E. Allen, L. Fin, J. Stark and A. Muir for help and assistance with tissue culture, the staff at the DNA sequencing service (School of Life Sciences, University of Dundee), the cloning, antibody and protein production teams within the MRC-PPU reagents and services (University of Dundee) coordinated by J. Hastie. We thank the staff at the Dundee Imaging Facility (School of Life Sciences, University of Dundee) for their invaluable help and advice throughout this project. We thank all members of the Sapkota lab for their highly appreciated experimental advice and/or discussions.

## Funding

KD is supported by an MRC Career Development Fellowship. GPS is supported by the U.K. Medical Research Council (grant numbers MC_UU_00018/6 and MC_UU_12016/3) and the pharmaceutical companies supporting the Division of Signal Transduction Therapy (DSTT) (Boehringer-Ingelheim, GlaxoSmithKline, Merck-Serono).

## Author contributions

KD performed the experiments, collected and analysed data, and contributed to the writing of the manuscript. TJM designed strategies and developed methods for the CRISPR/Cas9 gene editing, in addition to generating the cDNA constructs used in this study. GPS conceived the project, analysed data and contributed to the writing of the manuscript.

## SUPPLEMENTARY FIGURE LEGENDS

**Supplementary Figure 1:**
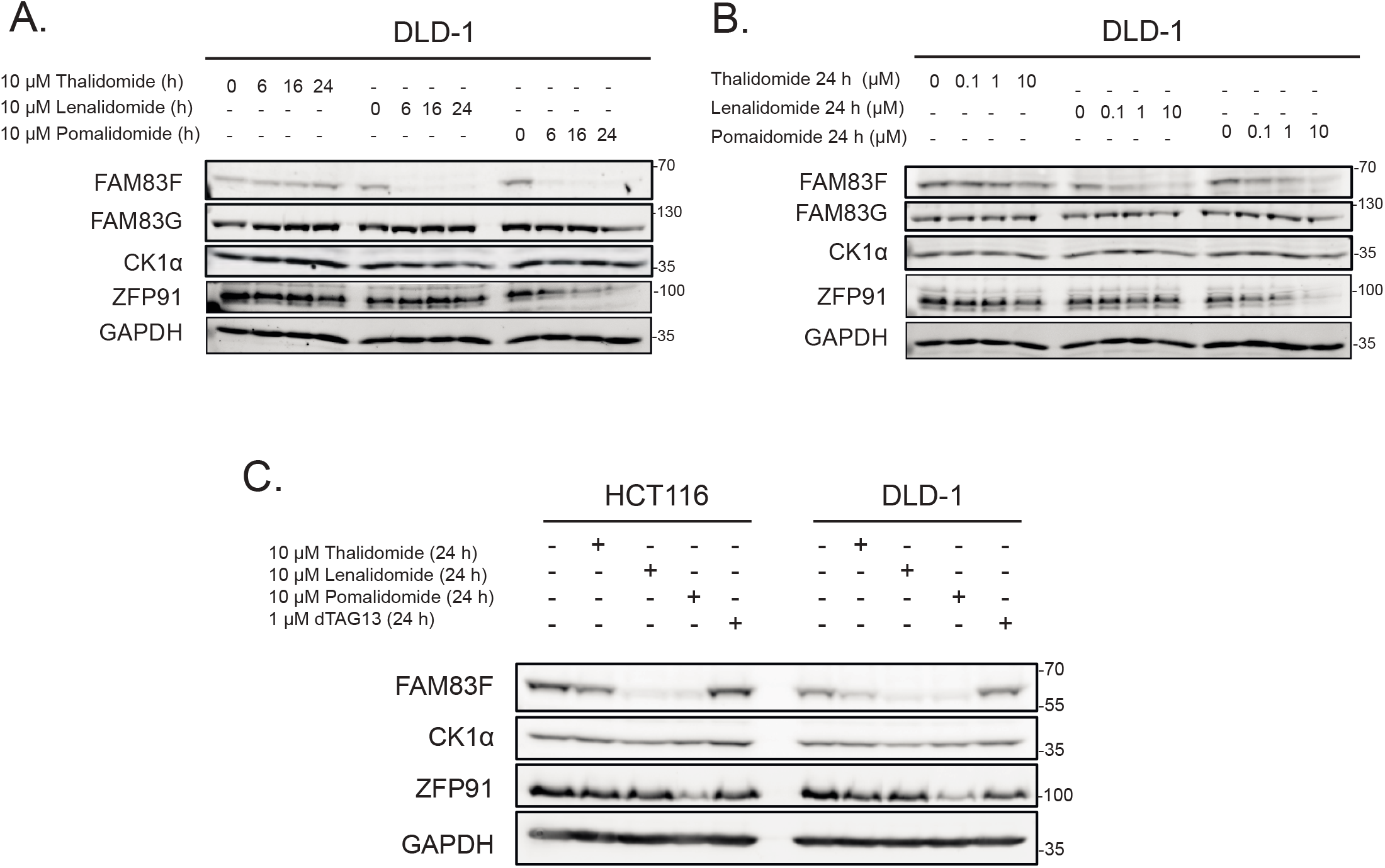
Characterisation of IMiD-induced FAM83F degradation in DLD-1 cells. **(A)** DLD-1 cell extracts treated with 10 μM IMiD compounds for various timepoints (6, 16 and 24 h) were resolved by SDS-PAGE and subjected to Western blotting with the indicated antibodies. **(B)** DLD-1 cell extracts treated with IMiD compounds for 24 h at various concentrations (0.1 μM, 1 μM and 10 μM) were resolved by SDS-PAGE and subjected to Western blotting with the indicated antibodies. **(C)** Cell lysates from HCT116 and DLD-1 cells treated with either IMiD compounds (10 μM 24 h) or dTAG13 (1 μM 24 h) were resolved by SDS-PAGE and subjected to Western blotting with the indicated antibodies.

**Supplementary Figure 2:**
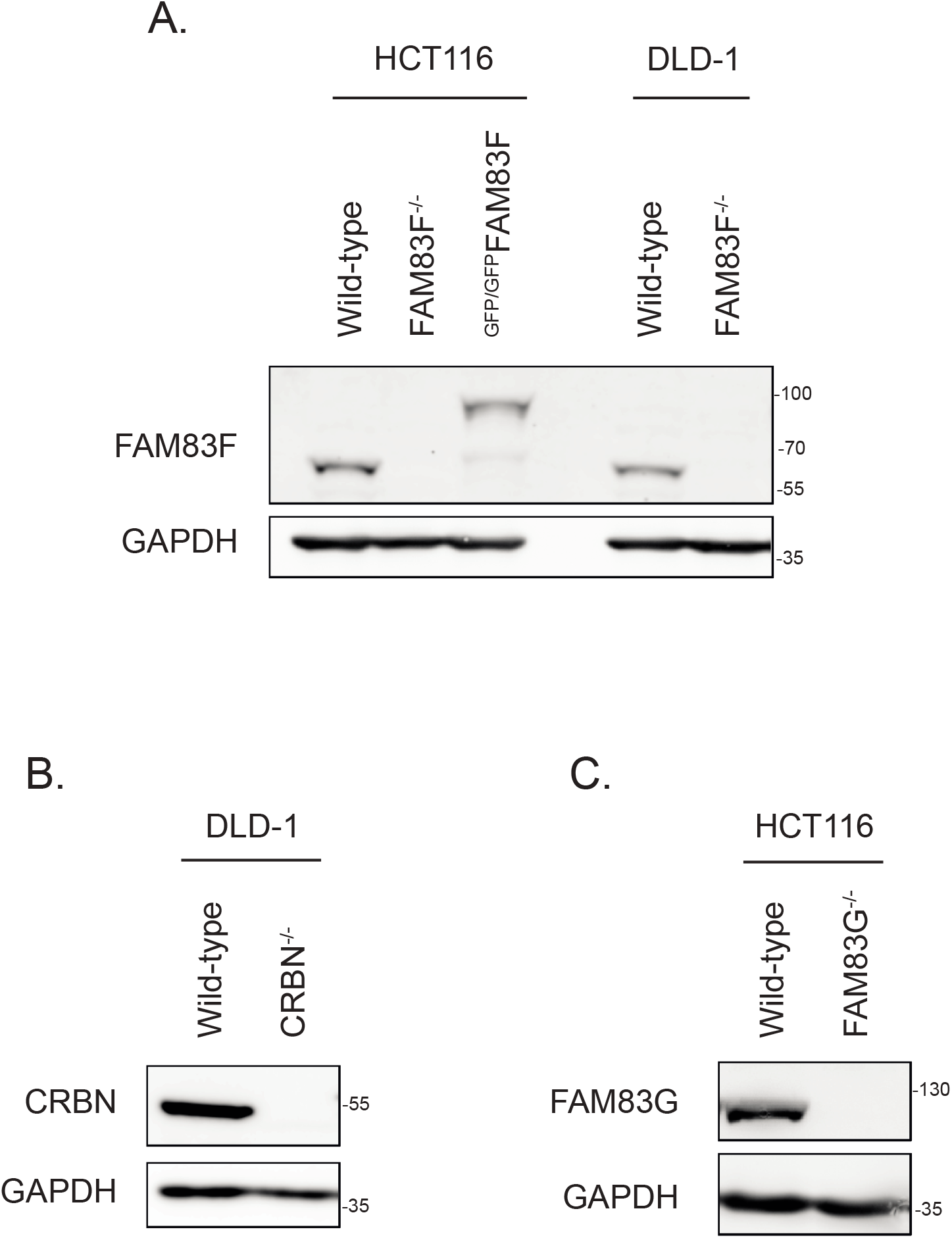
Western blot confirmation of CRISPR/Cas9 generated cell lines. **(A)** HCT116 (wild-type, FAM83F^-/-^ and ^GFP/GFP^FAM83F) and DLD-1 (wild-type and FAM83F^-/-^) cell extracts were resolved by SDS-PAGE and subjected to Western blotting with indicated antibodies. **(B)** DLD-1 wild-type and CRBN^-/-^ cell extracts were resolved by SDS-PAGE and subjected to Western blotting with indicated antibodies. **(C)** HCT116 wild-type and FAM83G^-/-^ cell extracts were resolved by SDS-PAGE and subjected to Western blotting with the indicated antibodies.

**Supplementary Figure 3:**
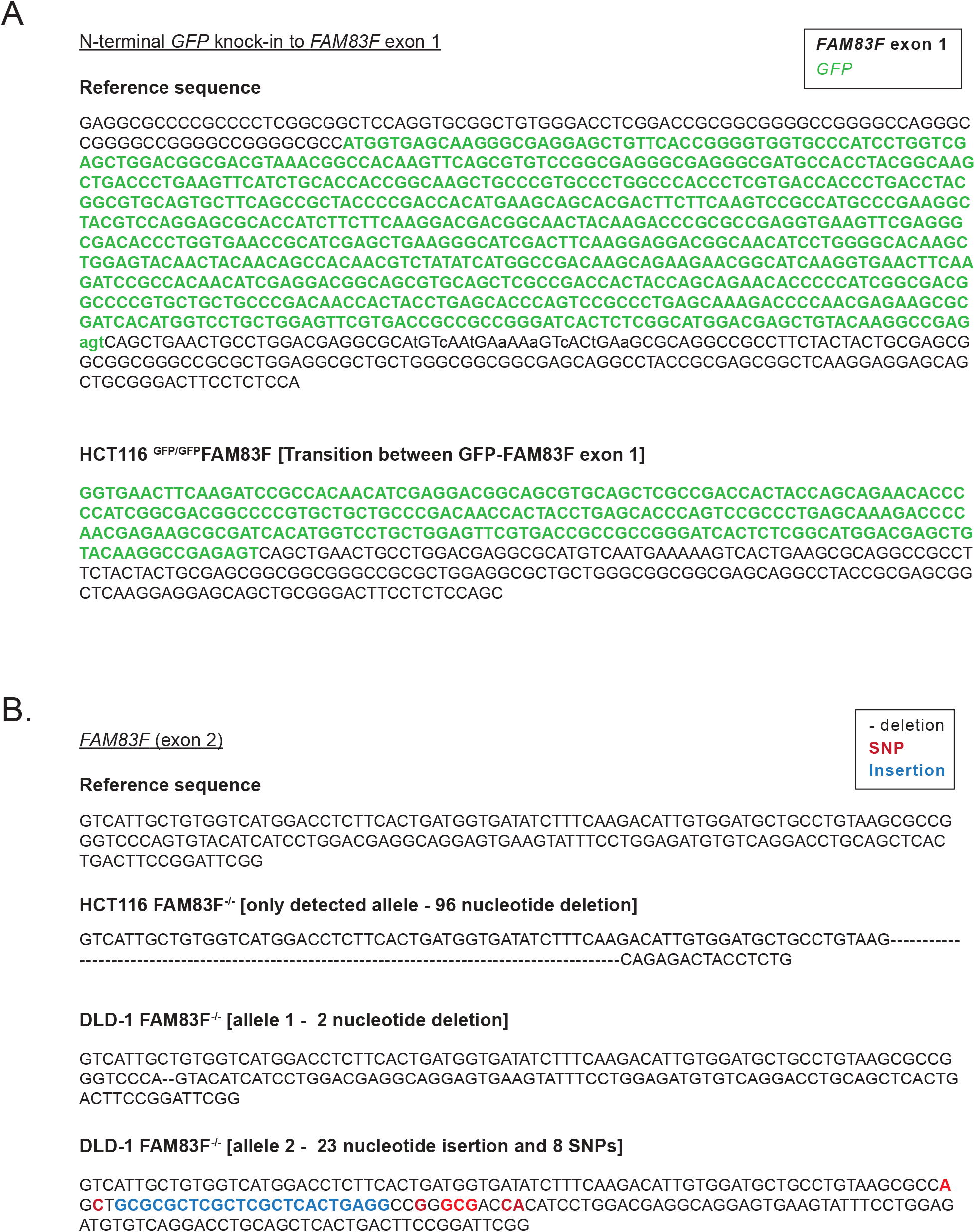
Sequencing confirmation of CRISPR/Cas9 generated FAM83F cell lines. **(A)** Reference DNA sequence of the CRISPR/Cas9 strategy to knock-in a *GFP* sequence into the N-terminus of the *FAM83F* locus and confirmation sequencing of the transition zone between *GFP* and *FAM83F* exon 1 in the HCT116 ^GFP/GFP^FAM83F cell lines. **(B)** Reference DNA sequence of *FAM83F* exon 2 and sequencing confirming alterations in the HCT116 FAM83F^-/-^ and DLD-1 FAM83F^-/-^ cell lines. These nucleotide changes predict a large deletion in the FAM83F protein sequence in HCT116 FAM83F^-/-^ cells, and the presence of premature stop codons in both detected FAM83F alleles for DLD-1 FAM83F^-/-^ cells.

**Supplementary Figure 4:**
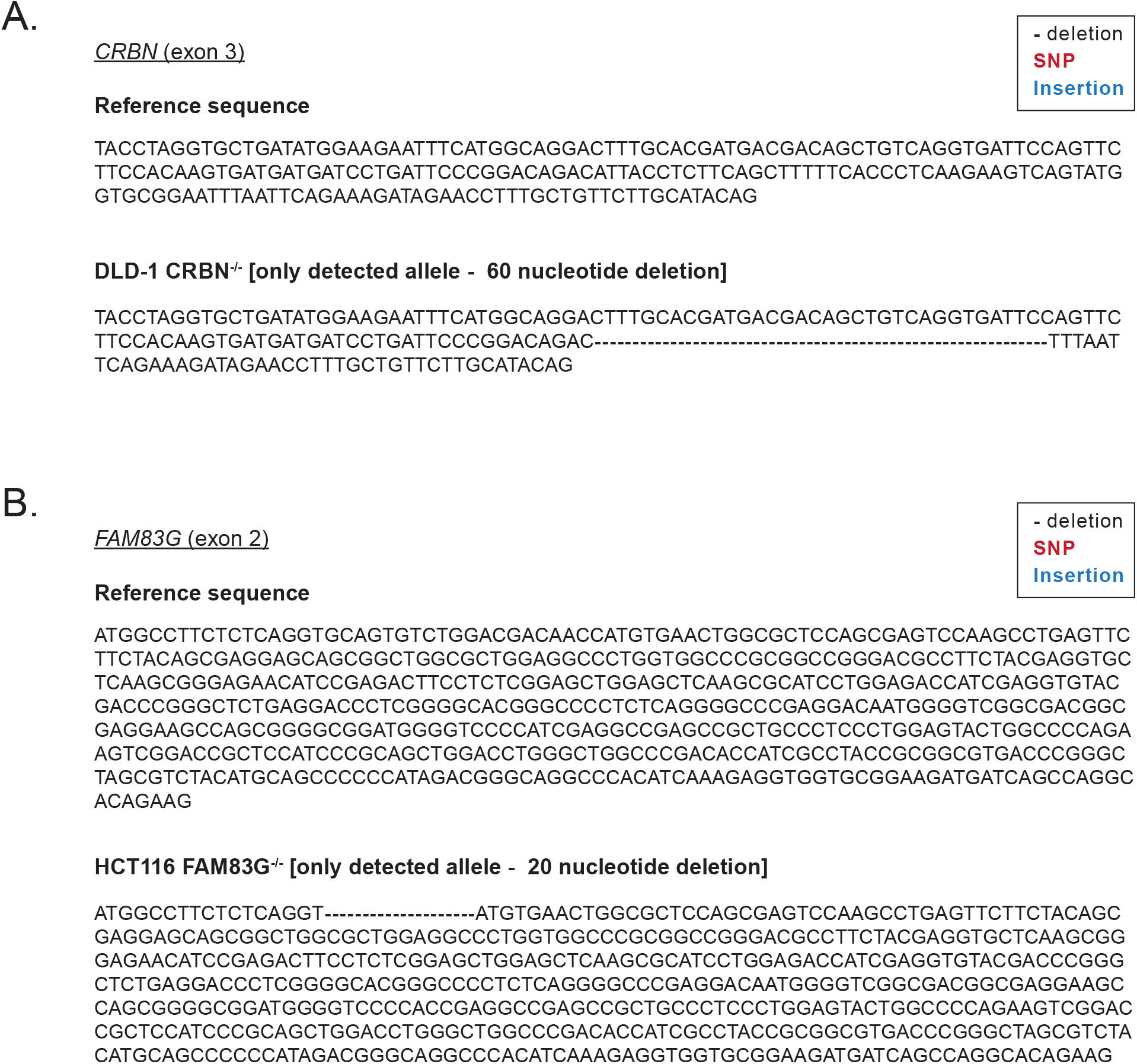
Sequencing confirmation of CRISPR/Cas9 generated CRBN and FAM83G cell lines. **(A)** Reference DNA sequence of *CRBN* exon 3 and DNA sequencing confirming alterations in the DLD-1 CRBN^-/-^ cell line. The nucleotide deletion predicts a deletion in the CRBN protein sequence for DLD-1 CRBN^-/-^ cells. **(B)** Reference DNA sequence of *FAM83G* exon 2 and sequencing confirming alterations in HCT116 FAM83G^-/-^ cell lines. The nucleotide deletion predicts a premature stop codon in the *FAM83G* locus in the HCT116 FAM83G^-/-^ cells.

**Supplementary Figure 5:**
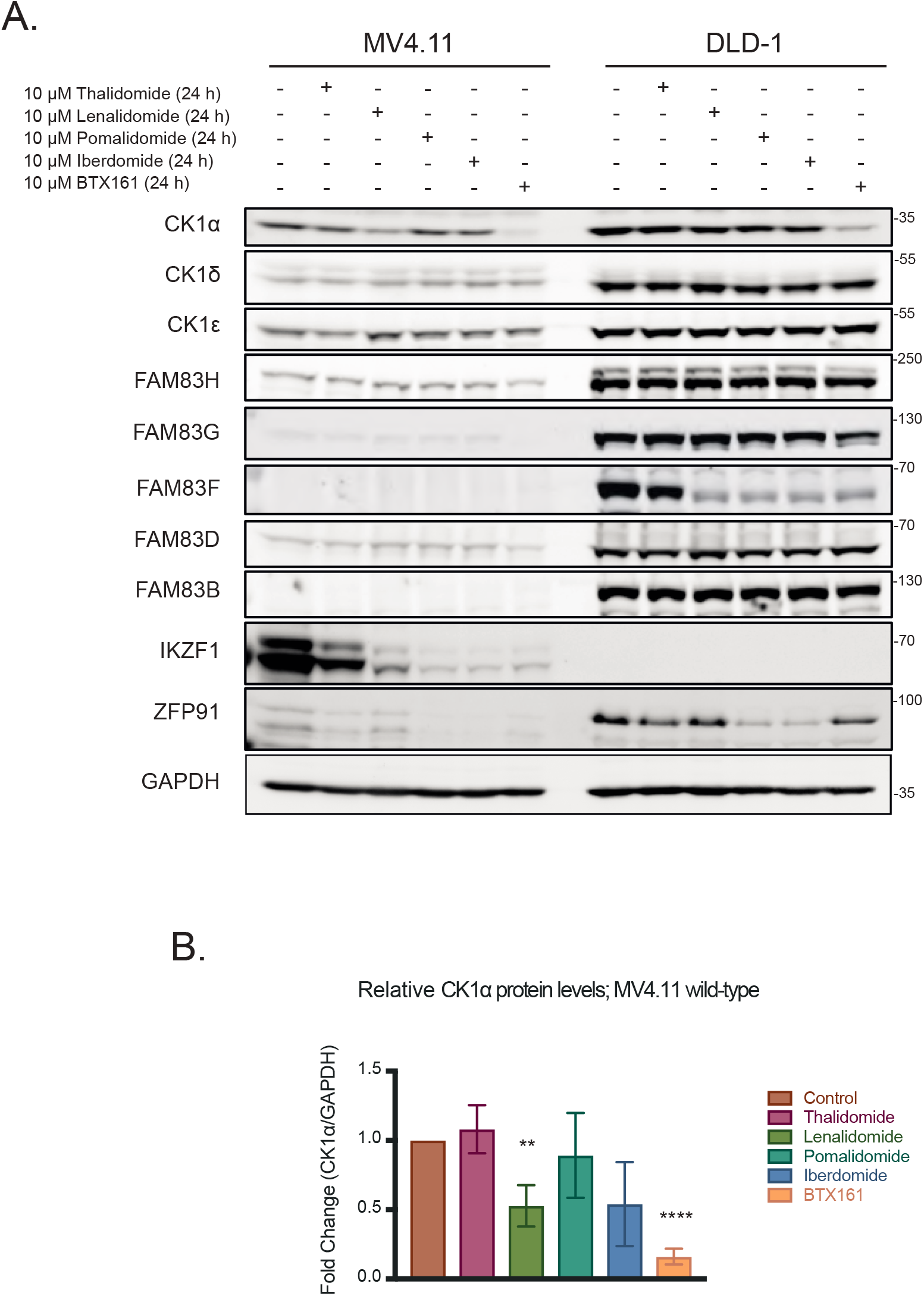
Endogenous FAM83 protein expression in MV4.11 cells. **(A)** MV4.11 and DLD-1 cell extracts treated with IMiD compounds (10 μM 24 h) were resolved by SDS-PAGE and subjected to Western blotting with the indicated antibodies. **(B)** Densitometry of CK1α protein abundance in MV4.11 cells treated with IMiD compounds (10 μM 24 h). CK1α protein abundance was normalised to GAPDH protein abundance and represented as a fold change compared to untreated cells. Data represents four biological replicates and bar graph represents mean ± standard error. Statistical analysis was performed using a students unpaired t-test and by comparing fold-change between untreated cells and cells treated with IMiD compounds (10 μM 24 h). Statistically significant p-values are denoted as asterisks (**** <0.0001, *** <0.001, ** <0.01, * <0.05).

**Supplementary Figure 6:**
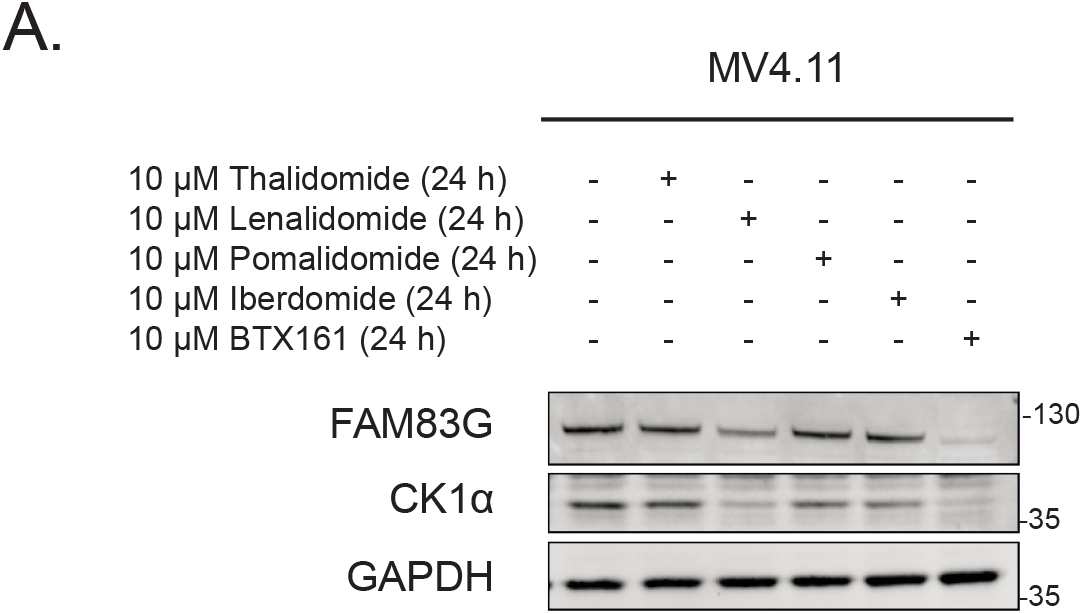
FAM83G degradation in MV4.11 cells. **(A)** MV4.11 cell extracts treated with 10 μM IMiD compounds for 24 h were resolved by SDS-PAGE and subjected to Western blotting with the indicated antibodies.

